# Convolutional neural network models describe the encoding subspace of local circuits in auditory cortex

**DOI:** 10.1101/2024.11.07.622384

**Authors:** Jereme C. Wingert, Satyabrata Parida, Sam Norman-Haignere, Stephen V. David

## Abstract

Convolutional neural networks (CNNs) provide powerful models of neural sensory encoding, but their complexity makes it difficult to discern computations that support their performance. To address this limitation, we developed a linear-nonlinear subspace model that identifies the most informative sensory dimensions captured by a CNN. A CNN was trained on single-neuron data recorded from auditory cortex of ferrets during presentation of a large natural sound set. Each neuron’s linear tuning subspace was computed by applying dimensionality reduction to the gradient of CNN output relative to input. Subspace projections were combined nonlinearly to predict neural activity. The resulting model was functionally equivalent to the CNN. Analysis of trained models showed that responses of local neural populations sparsely tiled a shared stimulus subspace. Encoding properties also differed between cell types and layers, reflecting their position in the cortical circuit. More generally, these results establish a framework for interpreting deep learning-based encoding models.

**Significance statement:** Auditory cortex mediates the representation and discrimination of complex sound features. Many models have been proposed for cortical sound encoding, varying in their generality, interpretability, and ease of fitting. It has been difficult to determine if and what different functional properties are captured by different models. This study shows that two families of encoding models, convolutional neural networks (CNNs) and tuning subspace models, account for the same functional properties, providing an important analytical link between accurate models that are easy to fit (CNNs) and models that are straightforward to interpret (tuning subspace).

## Introduction

Auditory perception requires parsing and grouping statistical regularities in the complex and highly variable sounds encountered in everyday environments. Numerous spectrotemporal encoding models have been proposed to describe the computations performed by neurons in the auditory system, and models with increasing complexity have been shown to account for sound-evoked activity with greater accuracy ^1–4^. Since the advent of deep learning methods, a number of studies have reported striking parallels between artificial networks fit to solve complex sensory problems, like face or speech recognition, and biological sensory representations, suggesting that these methods may also be used to model neural sensory processing directly ^5–7^. Consistent with this prediction, convolutional neural networks (CNNs) fit directly using data from the visual and auditory cortex perform substantially better than classical linear-nonlinear spectrotemporal receptive field (LN) models and variants of the LN model^8–11^.

Sensory encoding models that successfully predict time-varying neural activity can, in theory, be analyzed to determine what computations support their performance ^12^. A sufficiently accurate model of cortex should reflect local circuit properties, like dendritic integration and recurrent connectivity ^13,14^. However, developing encoding models that are both accurate and interpretable has proven challenging. The high prediction accuracy of CNN-based encoding models has been established, but their complexity and large number of free parameters has made it difficult to determine what nonlinear computations account for their performance. Given this problem, a concern about deep learning models is that they solve problems in ways that are not biological and thus their value to understanding biological computations may be limited ^15^. Being able to understand the key computations performed by a CNN can help us generate hypotheses about neural computations, which can then be tested directly against the biological network ^16,17^.

Subspace, or multifilter, models are a family of encoding models, derived from the LN modeling framework, that strike a balance between accuracy and interpretability. Subspace models define neural responses as a nonlinear function of the output of a relatively small number of spatiotemporal or spectrotemporal filters ^2,18,19^. The set of filters constrains the stimulus subspace that modulates a neural signal and can be easily visualized. Among other computations, subspace models can account for wide-ranging neurophysiological phenomena, such as gain control, and logical computations, such as “AND” gates, which cannot be captured by the LN model ^19–21^. While theoretically powerful and relatively easy to interpret, subspace models have proven difficult to fit, especially with natural stimuli, which contain high-order statistical correlations that present challenges to model fitting algorithms. Thus, studies that fit subspace models typically utilize simpler synthetic stimuli ^2,22,23^. Studies that have applied subspace methods to natural stimulus data have been constrained to small numbers of subspace dimensions ^24,25^, and published subspace analysis methods do not transfer readily to natural stimulus datasets ^10^.

Here we tested the hypothesis that a large, multi-layer CNN can be flattened into a low-dimensional encoding subspace, while maintaining high prediction accuracy. We used microelectrode arrays to record the activity of multiple single units across cortical columns of primary and secondary auditory cortex (AC) of awake ferrets during presentation of a large and diverse natural sound stimulus set. After fitting CNN models to neural data, we measured the tuning subspace of each neuron by a high-dimensional local linear approximation of the CNN followed by dimensionality reduction. A new encoding model based on a low (3-13) dimensional tuning subspace predicted AC neural activity nearly as accurately as the full CNN and revealed functional differences between neuronal cell types within a cortical column. Thus, subspace models derived from the CNN provide interpretable characterizations nonlinear tuning properties across cortical populations.

## Results

### A flattened convolutional neural network identifies the tuning subspace for auditory neurons

We developed a method to transform a deep and complex convolutional neural network (CNN) model into an interpretable subspace filter model of neural sound encoding. Single-unit neural activity was recorded across layers of auditory cortex of awake, passively listening ferrets using linear microelectrode arrays ^26,27^. Recordings were performed in primary (A1, 53 sites) and non-primary (PEG, 14 sites) auditory cortex (2874 units, 3-123 units/site, mean 43 units/site, 4 animals). To characterize the sound encoding properties of each neuron, data was collected during presentation of natural sound stimuli. The performance of encoding models is typically limited by the diversity of stimuli used to measure tuning ^12^. To maximize stimulus sampling diversity and match neural integration dynamics in the ferret ^28^, sequences of brief natural sound segments (duration 50-800 ms, mean 129 ms) were presented in rapid succession with occasional silent gaps (0.2-2 sec). The sound segments were drawn from a large natural sound library and sampled to maximize the diversity of spectrotemporal patterns presented. A single experiment presented approximately 6200-56000 unique sound segments (900-8100 sec) for model fitting.

A four-layer convolutional neural network (CNN) was used to model the functional relationship between the stimulus spectrogram and the time-varying neural response (Fig. 1A, Pennington and David, 2023). To leverage statistical power for model fitting, a population architecture was used, in which the first three model layers were shared across neurons within a recording site, and the final dense layer weighted activity from the final shared layer to predict the activity of individual neurons ^10^. The unique stimuli used for model fitting varied across recording sites. Model performance was evaluated using a held-out test set (six 18-sec segments), which was fixed across all experiments and presented 10-20 times on trials interleaved with the fit stimuli. Consistent with previous studies ^9,10^, the CNN model predicted time-varying neural activity in the test set more accurately than a linear-nonlinear spectrotemporal receptive field (LN) model (Fig. 1B-C, Fig. 2B, n=2337/2874 sound-responsive units, p<0.05 above-chance prediction for either model, permutation test corrected for multiple comparisons).

**Figure 1.**
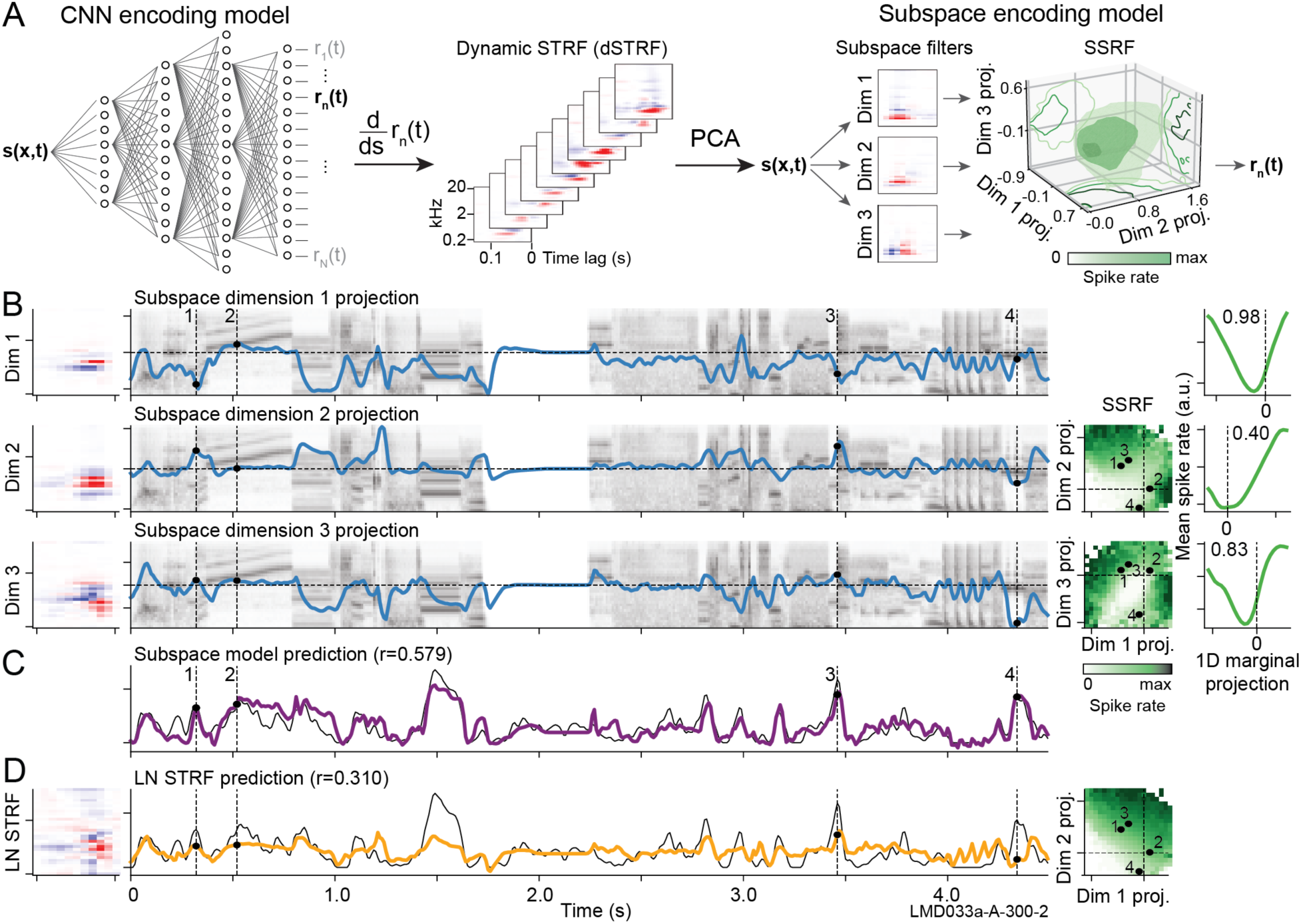
Mapping from convolutional neural network (CNN) to subspace encoding models. **A.** Population CNN model predicts time-varying neural activity recorded at a single site during presentation of a natural sound library. A dynamic STRF (dSTRF) is the collection of locally linear approximations of the CNN, computed as the gradient of the CNN response relative to the stimulus spectrogram. The neuron’s tuning subspace is estimated by principal components analysis (PCA) of the dSTRF across all timepoints. The subspace receptive field (SSRF) is then the mean response to stimuli projecting to each point in the tuning subspace. **B.** Subspace filters (left) are convolved with the stimulus spectrogram (second column, gray shading) to produce a projection into each dimension (3/7 highest-variance filters shown). The projection defines a point in the tuning space at each moment in time, and the SSRF indicates the predicted response at that time (third column, green shading). In this example, the SSRFs contain two distinct patches that produce a strong response (dark green, upper left and lower right in dimension 1 vs. 2 heatmap). Numbers indicate example timepoints (dashed vertical lines at left). Averages across x or y axes of the SSRF show the mean response as a function of the projection onto individual tuning dimensions (fourth column). In this example, marginal response functions are nonlinear and nonmonotonic, with strong responses for both positive and negative projections. Dashed vertical lines indicate zero stimulus (silence), which is not necessarily aligned to the low point of the tuning curves. **C.** Subspace model prediction (purple, aggregated across all significant filters) overlaid with the actual peri-stimulus time histogram (PSTH) response (gray, r=0.579). **D.** LN model fit for the same neuron (left), LN model prediction (orange) overlaid with actual PSTH (middle, r=0.310), and average LN model prediction for stimulus projections onto the first two dimensions of the subspace model (right). LN tuning is constrained to a one-dimensional subspace, which cannot capture the complex tuning of the SSRF in A.

**Figure 2.**
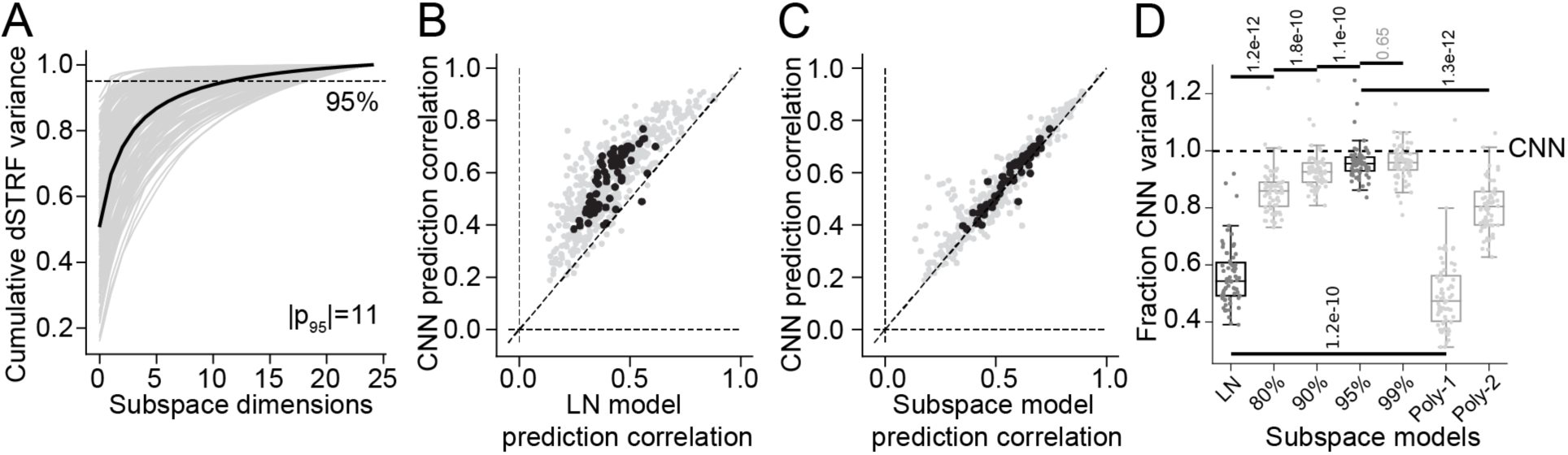
Equivalent performance of CNN and subspace encoding models. **A.** Thin lines show cumulative dSTRF variance explained as a function of principal component count for each neuron (n=2337 AC units). Thick line represents mean. The subspace encoding model used dimensions accounting for 95% of dSTRF variance (mean 11 dimensions). **B.** Scatter plot compares prediction correlation (noise-corrected correlation coefficient) between the LN and CNN encoding models. Gray dots indicate prediction correlation of single units, and black dots show the median for each recording site (median across all units 0.416 vs. 0.600, p<1.0e-10, Wilcoxon signed-rank test). **C.** Scatter plot comparing subspace (SS) versus CNN model performance, plotted as in C (median 0.584 vs. 0.600, p=1.0e-7). **D.** Box plots compare model performance, as a fraction of response variance explained (squared correlation coefficient) relative to the full CNN encoding model (dashed line). Dots indicate site medians (n=67, models from B and C highlighted in dark gray). Subspace models were tested with variable number of PCA dimensions used to define inputs (80-99% of dSTRF variance). Polynomial models (right) constrained the mapping from subspace to neural response to be linear (Poly-1) or a second order polynomial (Poly-2). For this dataset, prediction accuracy was maximal for the 95% subspace model and did not increase significantly for the 99% subspace model. Black bars highlight performance comparisons between pairs of models (Wilcoxon signed-rank test, numbers indicate exact *p* values).

While the CNN-based auditory encoding model can account for neural response dynamics more accurately than a traditional LN model, the multiple layers of nonlinear processing make it difficult to understand what computations account for the increased predictive power. We developed a method to “flatten” the layers of a deep network into a compact set of transformations that can be visualized more readily. Previous work has shown that a CNN-based encoding model can be visualized with the dynamic STRF (dSTRF), the derivative of the model response relative to the input stimulus at each point in time (Eq. 5) ^9^. Analytically, this method is similar to saliency analysis sometimes used to understand deep learning models ^29^. dSTRF analysis produces a large collection of spectrotemporal filters (Figs. 1A), which provide a piecewise linear approximation of the stimulus-response relationship, with each dSTRF centered around the spectrotemporal input at a different point in time. dSTRFs for a single neuron often vary substantially but show some regular patterns in their structure (Fig. S1). For a compact visualization of dSTRF variability, we performed dimensionality reduction using principal components analysis (PCA) on the collection of dSTRFs for each neuron. For most A1 neurons, 3-16 principal components (mean 11) accounted for most of dSTRF variance (>95%), and 3 components accounted for a mean of 81% of dSTRF variance (Fig. 2A). The components measured for a single neuron often shared a best frequency but differed in spectral and/or temporal modulation tuning (examples in Figs. 1B, S2-S5).

Previous studies of sensory encoding have proposed that neural representation can be described by a subspace, or multifilter, encoding model ^2,19,30–32^. In this family of models, each neuron is characterized by one or more linear filters, and sensory responses are predicted entirely from the projection of the stimulus into this low-dimensional subspace. In the case of one filter, the subspace model reduces to the LN model. The large collection of filters in the dSTRF describes the gradient of model outputs over the full set of fit stimuli. We surmised, thus, that the principal components of the dSTRF defined a tuning subspace analogous to that of multifilter models. To visualize how the subspace can account for sound encoding, we computed the largest principal components of the dSTRF for each neuron. We projected the stimulus into this subspace and computed the average neural response across stimuli at each point in this representation (Figs. 2, S2). *Subspace receptive fields* (*SSRFs*) show the average response to the stimulus projected jointly onto a pair of dimensions (Fig. 1B, second column from right), and subspace tuning curves show the average response to stimuli varying along a single subspace dimension (Fig. 1B, right column). Analogous to receptive fields defined with respect to receptors in peripheral sensory areas, the SSRF indicates the region of the tuning subspace that evokes a spiking response. The SSRF can be nonmonotonic and have multiple peaks. In contrast, predictions of the LN model projected into the same subspace are constrained to lie in a one-dimensional subspace consistent with a single linear filter instantiated in the LN model (Fig. 1C).

### Subspace encoding model is functionally equivalent to the CNN

The spectrotemporal tuning subspace provides a flattened neural network model. A CNN may contain many layers of many units, but the subspace tuning model describes the underlying computations as convolution by a single layer of spectrotemporal filters followed by a nonlinear combination of those filter projections, reflected in the SSRF. The SSRF provides a lookup table for the neural response at each stimulus timepoint, thus providing a compact visualization of the nonlinear computations that define a neuron’s encoding properties (Fig. 1).

While the subspace model provides a visualization of tuning captured by the CNN, it need not account for neural responses as accurately as the full model. To determine how well the subspace representation accounts for time-varying neural responses, we fit a new model (Fig. 2A), where neural responses were predicted using only the projection into the tuning subspace. A small neural network with three fully connected layers was used to fit the receptive field within the subspace (SSRF, Eq. 7). The subspace model was able predict neural responses with nearly the same accuracy as the full CNN (median r=0.585 vs. 0.600, p=1.0e-7, Wilcoxon signed-rank test, Fig. 2C). Overall, the subspace model accounted for an average of 0.954 of the response variance explained by the CNN (Fig. 2D). A direct comparison of time-varying neural activity predicted by the two models showed strong correlation, indicating that the two models are functionally nearly equivalent, while equivalence was substantially lower between the CNN and LN models (Fig. S3A). Moreover, the shape of SSRFs fit for the subspace model were strongly correlated with the mean actual response to stimuli projected into the tuning subspace, indicating that tuning in the SSRF did in fact account for sound features that modulated the neural response (Fig. S4).

The dimensionality of the tuning subspace is not fixed *a priori*, and some criterion must be specified for how many PCA dimensions to include in the subspace (Fig. S5). We compared performance for models using variable numbers of subspace dimensions, determined by the fraction of dSTRF variance explained. Median cross-validation performance improved for models including filters accounting for up to 95% of dSTRF variance. There was no additional improvement for higher dimensionality, indicating that the additional dimensions were dominated by noise in the CNN fit (Fig. 2D). When results were broken down by cortical subregion, similar performance was observed for the subspace model in both areas (A1: 0.955, PEG: 0.946 median fraction of CNN variance explained). Thus, the relatively small number of subspace dimensions captures the majority of the functional response characteristics of the full CNN.

The tuning subspace describes the set of spectrotemporal patterns that influence a neuron’s activity, but it does not constrain the nonlinear computations performed within that space. If a neuron’s response is simply a linear weighted sum of the stimulus subspace projection, it should behave like a single linear filter, as in the LN model (Fig. 2B). Indeed, a model constrained to use a linear combination of subspace projections (Poly-1) performs similarly to the LN model (0.491 vs. 0.564 median fraction CNN variance explained, Fig 2D). The slightly worse performance likely reflects the absence of a sigmoid nonlinearity in the Poly-1 model. Previous studies implementing spike-triggered covariance have modeled subspace tuning with a second-order polynomial ^19,21^. A second-order model in the tuning subspace performs better than the LN model, but worse than the full CNN or full subspace model (Poly-2, 0.829 relative variance, Fig. 2D). Thus, while second-order interactions account for some of the subspace tuning, higher-order nonlinearities are required to fully describe the SSRF.

Previous attempts to fit subspace models directly to neural data, without an intervening CNN-based model, have had mixed results and generally required carefully tuned hyperparameters ^2,10,25,30,32^. For the current data, subspace models trained directly on the neural data performed similarly to but not as accurately as those estimated using the CNN-based analysis (Fig. S3B-D). The CNN-based method also produced less noisy subspace filters, and measures of subspace tuning from the CNN were also stable, even when only a relatively small number of dSTRFs were included in the PCA (Fig. S6A-D). The use of natural sounds produced more compact measures of the tuning subspace. When a new subspace model was constructed using an equivalent number of random noise stimuli to define subspace filters, the resultant model showed slightly lower prediction accuracy and required more dimensions (i.e., subspace filters, Fig. S6E-F). Thus, the natural sound-based model provides a compact and efficient characterization of natural sound encoding. This model may not generalize to arbitrary noise stimuli as accurately as a model fit with a larger synthetic stimulus set, but it is adequate for the natural stimuli used here.

### Neurons within a cortical column sparsely tile the local tuning subspace

The linear microelectrode arrays recorded the activity of multiple neurons across layers at the same location in cortex. Thus, the tuning of simultaneously recorded neurons within versus across recording sites provided insight into sound features represented topographically in AC. The most prominent feature in AC is tonotopy, smooth changes in neuronal best frequency that define AC subfields ^33,34^. In addition, topographic organization has been reported for tuning to multiple other features, including spectral bandwidth, temporal modulation rate, and binaural contrast ^35–37^. Within a single recording site, subspace filters showed broad similarity in tuning to stimulus frequency and bandwidth (examples in Fig. 3A, Fig. S5). However, more fine-grained tuning to spectral and temporal modulations differed across neurons.

**Figure 3.**
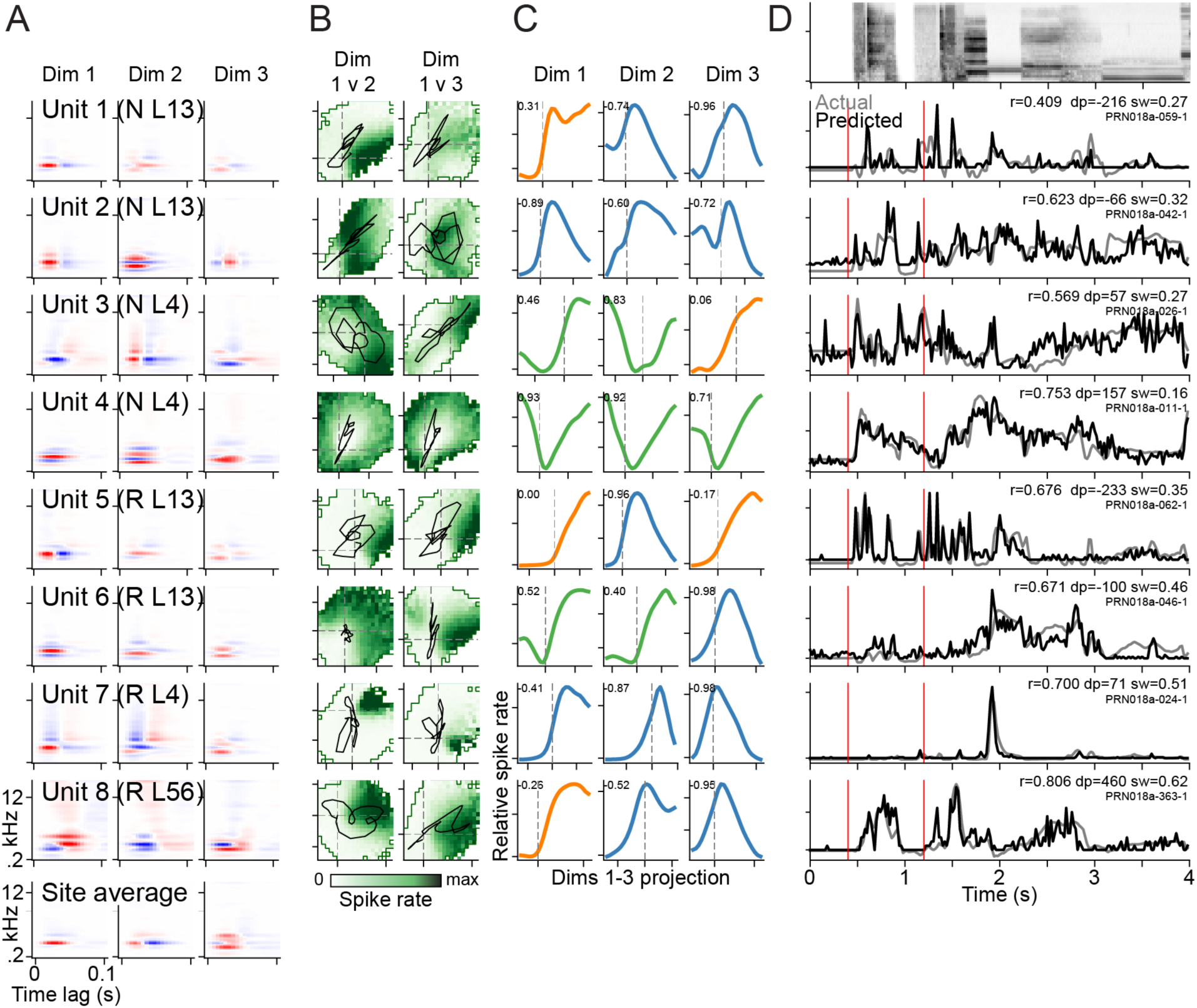
Diversity of subspace models within a single recording site. **A.** Three largest-variance subspace filters for eight example units, one per row. Labels for each unit indicate spike width (N: narrow, R: regular) and cortical depth (L13: layers 1-3, L4: layer 4, L56: layers 5-6). Bottom row shows site-wide subspace filters, computed from dSTRFs aggregated across all units. **B.** Two-dimensional subspace receptive fields (SSRFs) for dimensions 1 vs. 2 (left) and 1 vs. 3 (right), computed as the average predicted response to stimuli projecting to each point in the tuning subspace. Darker green indicates higher spike rate. Black lines show trajectory of stimuli in the subspace during 0.4-1.2 sec in panel D. Thin green lines outline the valid SSRF region, i.e., the area sampled by at least five occurrences of the stimulus used for model fitting. **C.** Tuning curves show marginal projections of the SSRF along individual subspace dimensions. Numbers indicate tuning symmetry index (TSI, Eq. 10), and color indicates TSI category (orange: asymmetric, blue: symmetric downward-facing, green: symmetric upward-facing). Dashed vertical lines indicate zero stimulus (silence), which is not necessarily aligned with the minima/maxima of symmetric curves. **D.** Top row shows 4-sec segment of spectrogram from the natural sound sequence used to test prediction accuracy. Each row below shows the predicted PSTH response (black) overlaid with the actual response (gray). Numbers indicate prediction correlation (*r*), microns below cortical layer 3/4 boundary (*dp*) and spike width in ms (*sw*).

Nonlinear tuning within the subspace also varied. The SSRF was sometimes confined to a small region, producing sparse responses to sounds, but other times showed broad tuning and responses to a wide range of stimuli (compare units 6 and 7, Fig. 3B-C). Thus, despite sharing a similar tuning subspace, PSTH responses differed substantially within the same recording site (Fig. 3D).

To quantify tuning similarity within a recording site, we compared the tuning subspace for each neuron to the shared subspace, computed for the entire recording site (Fig. 3A). The shared tuning subspace was computed by concatenating dSTRFs across all units in the same site. We performed the same PCA procedure as for single units, above, and the dimensions explaining the most dSTRF variance defined the stimulus subspace for the entire site (Fig. 3A, bottom row). We quantified similarity between the single neuron- and site-wide filters using a subspace similarity index (SSI, Eq. 9), the mean cosine similarity between all combinations of the four largest subspace filters, which accounted for an average of 85% of dSTRF variance across all recording sites (Fig. 1B). An SSI of 1 indicates a complete alignment of subspaces, and SSI of 0 indicates no alignment (i.e., orthogonality). Within-site SSI was contrasted against SSI computed between the neuron and the shared subspace for a different recording site. Because population CNN models were fit simultaneously for all neurons within a site, the shared model layers could bias subspace filters to be similar and, thus, bias SSI toward higher values. To control for this potential bias, the SSI comparison was restricted to experiments in which the same natural sounds were presented during recordings from 2-3 different sites (40 sites total). A single model was fit using data pooled across sites, and SSI was compared only between neurons in the same model fit. Thus, any bias toward similarity was balanced for within-site and between-site comparisons. The comparison revealed that SSI was greater within than between sites (median 0.55 vs. 0.35, p=2.5e-11, sign test, Fig. 4B). This difference was the same when fewer or more than four subspace dimensions were used to compute SSI (Fig. S7). Thus, neurons recorded from the same site tended to share a similar tuning subspace.

**Figure 4.**
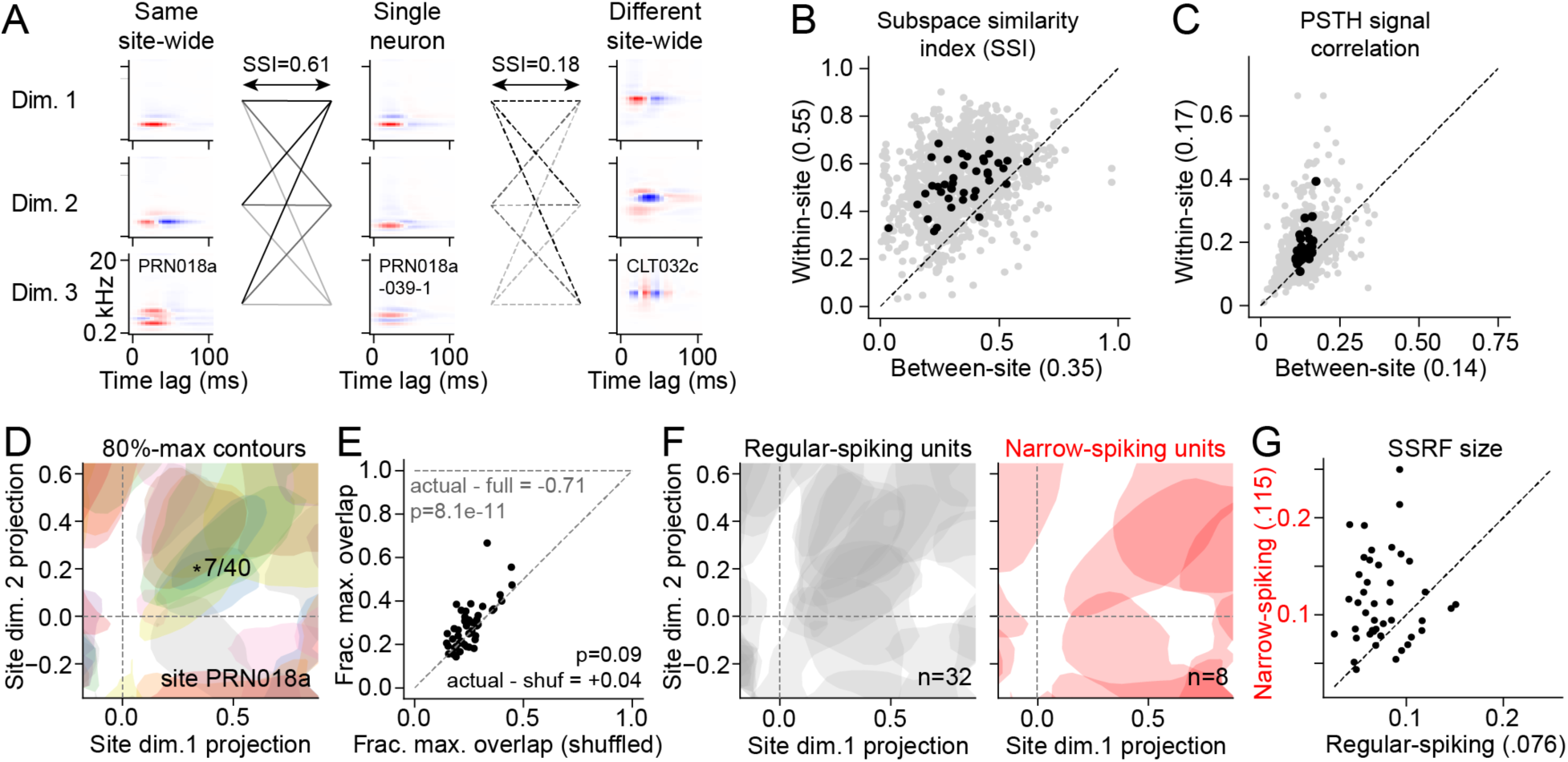
Sparse tiling of a shared tuning space within recording sites. **A.** Subspace similarity index (SSI) compares two tuning subspaces by computing the mean cosine similarity between all pairs of dimensions in the two subspaces. SSI was computed between each neuron and the site-wide average of the same site (within-site) and a different site presented with the same stimuli and fit simultaneously with the same population CNN (between-site). **B.** Scatter plot compares SSI for single units and site-wide averages between-site and within-site. Gray dots show individual units; black dots show mean for each recording site. SSI within-site is significantly greater than between-site (0.55 vs. 0.35, p=2.6e-11, sign test, n=39 recording sites). **C.** Scatter plot of correlation between predicted PSTH response (signal correlation) for pairs of units, using the validation dataset and plotted as in B. Correlation between predicted PSTHs was relatively low in all cases, but median correlation within-site was significantly greater than between-site (|0.17 vs. 0.14, p=3.6e-8, Wilcoxon signed-rank test). **D.** SSRF of each neuron (n=40, same site as Fig. 3) computed for dimensions 1 and 2 of the shared, site-wide tuning subspace. Filled contours show the high-activity SSRF where each unit reaches >80% of maximum firing rate. The full set of SSRFs tiles most of the subspace, and maximum overlap across the subspace is 7/40 (*). **E.** Scatter plot of fraction maximum overlapping SSRFs in the shared dimension 1 and 2 subspace for shuffled-versus actual high-activity SSRFs (n=39 sites). Fraction of maximum overlap was not significantly different from overlap in the shuffled SSRFs (p=0.09, Wilcoxon signed-rank test). **F.** High-activity SSRFs of regular-spiking units (n=32, left, gray) and narrow-spiking units (n=8, right, red), replotted for site in D. **G.** Scatter plot compares mean fraction of subspace area covered by SSRFs across each recording site, for regular-versus narrow-spiking neurons (n=39 sites, mean regular: 0.076, narrow: 0.115, p=4.4e-5, Wilcoxon signed-rank test).

Despite the similarity in tuning subspaces within a recording site (e.g., Fig. 3A), PSTH responses to the same stimuli varied substantially across neighboring neurons (Fig. 3D). To gain a clearer understanding of shared encoding properties within a recording site, we measured the signal correlation (correlation coefficient) between the PSTH predicted by the subspace model for pairs of neurons within and across sites (Fig. 4C, data matched to the SSI comparison). Despite the high SSI within site, average signal correlation was relatively low, although it was still higher within-site than between sites (median 0.17 versus 0.14, p=3.6e-10, sign test). Thus, despite spanning similar tuning spaces, pairs of neurons tended to produce very different time-varying activity.

The similarity of tuning subspaces within a recording site suggested that response properties of individual neurons could be projected into and compared within the shared, site-wide tuning space. For each neuron, we computed the SSRF in the site-wide tuning space. We defined the high-activity region of the SSRF as the area with >80% of its maximum value, initially focusing on the SSRF for the two largest subspace dimensions. High-activity SSRFs for individual neurons often spanned a relatively small area and were distributed across the tuning subspace. The substantial scatter of SSRFs within the subspace is consistent with the observation that PSTH responses differ substantially between units (Fig. 3D). Marginal tuning curves were often symmetric around non-zero stimulus values (Fig. 3C), explaining why many high-activity SSRFs were centered at locations away from the origin (intersection of dashed lines in 3B) in the subspace. This distribution suggests that tuning of a local population sparsely tiles the shared tuning subspace. An implication of sparse tiling is that high-activity regions of SSRFs for different neurons should not often overlap. To test this prediction, we measured the maximum overlap (i.e., intersection) of high-activity SSRF regions across an entire site. In an example site with 40 units, the maximum overlap was 7 (20% of units, * in Fig. 4D). If neurons all shared the same tuning, maximum overlap should be equal to the number of units. If tuning is random, the amount of overlap will depend on the size of the SSRFs. To test how closely the SSRFs resembled a random distribution, we shuffled the position of each SSRF and computed a maximum average overlap of 6 (Fig. S8). Thus, the number of neurons with simultaneous high activation for any stimulus was nearly the same as for a collection of randomly organized SSRFs. We observed the same result across subspace dimensions and recording sites (Fig. 4E, S8). Maximum overlap varied with the number of neurons per site, but on average it was only 4% greater than expected by chance and 71% less than full overlap, i.e., if all neurons shared the same SSRF. Thus, while neurons in a recording site tend to share a spectrotemporal tuning subspace, their responses are relatively uncorrelated, and their SSRFs tile the shared subspace.

SSRFs typically spanned a small region of the tuning subspace, but their size did vary (compare units 6 and 7 in Fig. 3). Previous studies have found that some inhibitory interneurons have broad sensory tuning ^38^, which would be consistent with larger SSRFs. To determine if we could explain variability in SSRF size by cell type, we used action potential spike width to classify units as putative excitatory (regular spiking) or inhibitory (narrow spiking) neurons ^39^. We measured peak-to-trough width of each spike and observed a bimodal distribution, delineating a boundary of 0.35 ms between regular- and narrow spiking groups (Fig. S9B ^40^). We then compared the area of high-activity SSRFs between these groups. While narrow-spiking neurons were less common, each group of neurons tiled the tuning subspace (Fig. 4F). Narrow spiking neurons also had consistently larger SSRFs than regular spiking neurons (Fig. 4G). Thus, in the same way that inhibitory neurons can have broader tuning to sound features like tone frequency ^41^, their SSRFs also span a larger area of the site-wide tuning subspace.

### Local subspace overlap depends on neuronal cell type and cortical depth

While neurons in the same recording site shared broadly similar tuning subspaces (Fig. 4B), there was still variability between neurons (Fig. 3A). We next asked what factors explain this within-site variability of tuning subspaces. The insertion of electrode arrays normal to the brain surface sampled activity across depths in the same cortical column. We used current source density and local field potential power profiles to determine depth, using standard methods (Fig. S9A, ^42,43^). We also classified units as regular spiking (putative excitatory) and narrow spiking (inhibitory), as described above (Fig. S9B). Model prediction accuracy was slightly higher in superficial cortical layers and for narrow spiking neurons (Fig. S9C). Average spike rate was also higher for narrow spiking neurons, consistent with previous reports of higher tonic firing rate in inhibitory neurons (Fig. S9E, ^14^).

A comparison of SSI between all pairs of neurons in an example recording site showed that subspace similarity varied with both cortical depth and spike width (Fig. 5A). SSI tended to be higher for narrow spiking neurons in superficial layers and lower for regular spiking neurons in deeper layers. This pattern is consistent with observations for individual neurons (e.g., compare similarity of Fig. 3A units 1-2 versus units 7-8).

**Figure 5.**
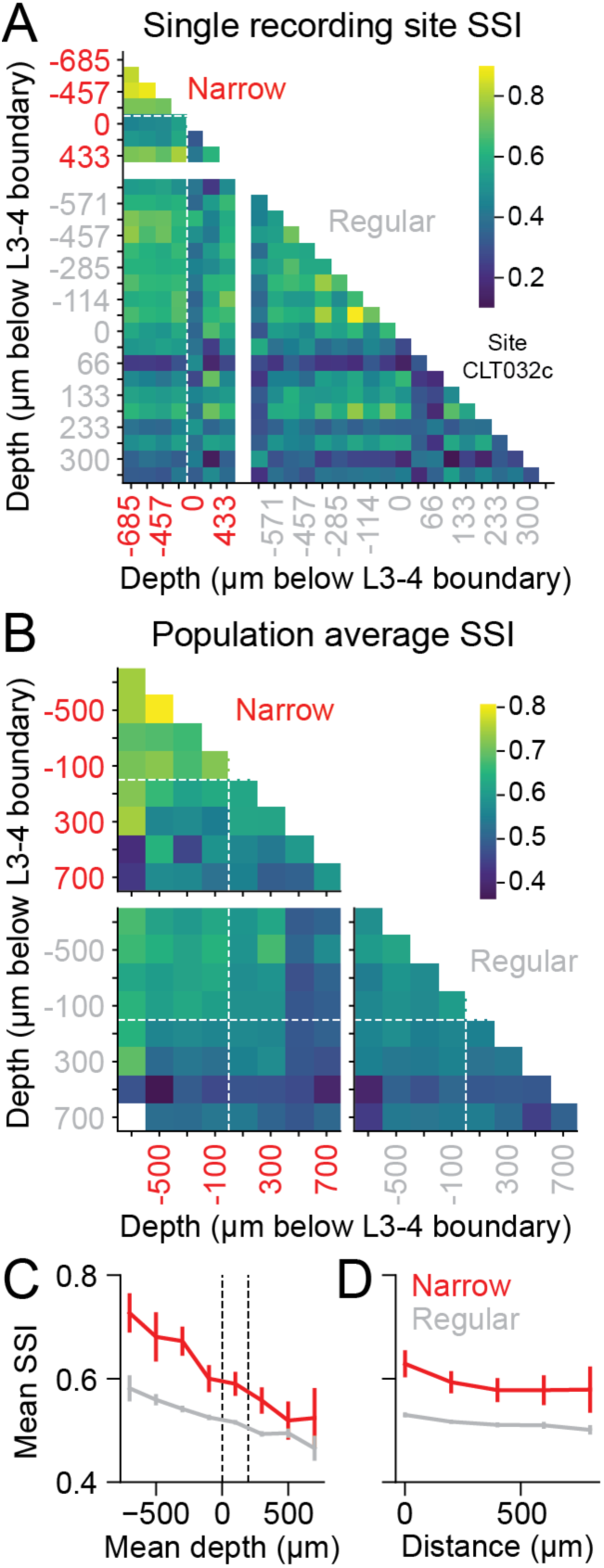
Subspace similarity within a local A1 population depends on neuronal cell type and cortical depth. **A.** Heatmap shows subspace similarity index (SSI) between each pair of single units from a single recording site, grouped by spike width (narrow, red; regular, gray) and then sorted by cortical depth. **B.** Mean SSI between pairs of neurons within recording site, grouped by spike width and 200 um bins, and averaged across all sites (narrow: n=205 units, regular: n=1459 units, 42140 pairs total, Fig. S9). **C.** Mean SSI by cortical depth (x axis) and spike width (color), for each unit paired (i.e., average across each depth in upper and lower triangles in B). Both cortical depth and spike width significantly predicted SSI (Table S1). Error bars indicate 2 SEM and are larger for narrow spiking neurons because of their smaller sample size. Vertical lines indicate boundaries between cortical layers 3-4 and 4-5. **D.** Mean SSI as a function of distance (difference in depth) between the neuron pair (x axis) and spike width (color), plotted as in C.

We averaged effects of cell type on SSI across recording sites for pairs of neurons grouped by their cortical depth (200-μm bins) and spike width (narrow vs. regular). Consistent with the example site, average SSI was higher between pairs of narrow spiking neurons in superficial layers (Fig 5B-C). This tendency toward higher SSI in superficial layers was driven primarily by the average depth of the units being compared. SSI decreased slightly for neurons separated by a large difference in depth, but this effect was weaker than the dependence on average depth (Fig. 5D). We quantified the dependence of SSI on cell type with a linear mixed-effects model (Eq. 11). For neurons with less reliable auditory responses, we expected the population model architecture to impose tuning properties that resembled the site average and thus inflate SSI between pairs of noisy neurons. To control for this potential bias, prediction correlation and mean spike rate (both correlated with response reliability, ^44^) were included as regressors in the model. Animal identity (*n*=4) was also included as a random effect to account for between-subject variability.

The full mixed-effects model revealed that both mean depth (averaged between neurons) and spike width (narrow-narrow, narrow-regular, regular-regular categories) significantly predicted SSI in A1, confirming the greater SSI between pairs of superficial and narrow spiking neurons (depth, χ^2^=566.6, p<2e-16; spike width, χ^2^=495.8, p<2e-16; width x depth, χ^2^=52.8, p=3.5e-12, n=42140 pairs, see Table S1 for detailed results). Model prediction correlation and neurons’ mean spike rate also had significant effects (prediction, χ^2^=1052.5, p<2e-16; rate, χ^2^=290.9, p<2e-16). These prediction accuracy and spike rate effects are likely to be independent of cell type effects, since depth and spike width showed relationships to these variables distinct from their relationship to SSI (compare Fig. 5C and Figs. S9C-D). As a particular example of this dissociation, narrow spiking neurons had both higher SSI and higher prediction accuracy, but prediction accuracy is negatively correlated with SSI on average. Data from PEG also showed the effects of cortical depth and spike width, although effect size was slightly weaker, possibly reflecting the smaller sample size of this dataset (Table S1).

### Diversity of nonlinear responses within the tuning subspace

Inspection of the subspace models shows a diversity of SSRFs across neurons, varying in size and shape (Fig. 3). These differences could reflect distinct elements of the underlying cortical circuit. As demonstrated above, larger SSRFs are associated with narrow-spiking, putative inhibitory neurons (Fig. 4F-G). To further quantify SSRF properties, a marginal tuning curve was calculated for each subspace dimension by projecting the stimulus into that single dimension and measuring the mean response for each value of that projection (Fig. 3C). Marginal tuning curves varied in their degree of symmetry ^45^, ranging from completely asymmetric (as would be expected for the linear-nonlinear model) to completely symmetric (consistent with a second-order polynomial model, examples in Fig. 3C). Among symmetric tuning curves, the majority were downward facing, producing high firing range over a relatively small, bounded range of inputs and a decrease in firing rate at large positive and negative values. In addition, a minority of tuning curves were upward facing, producing high firing rate for large positive and negative inputs. Inspection of example neurons showed that the downward versus upward orientation of the symmetric tuning curves had distinct impact on the time-varying neural response. Neurons with downward-facing tuning curves tended to respond sparsely to a small number of stimuli (e.g., Fig 3, unit 7), while neurons with upward facing tuning curves tended to respond to many stimuli (e.g., Fig. 3, unit 4). Thus, the properties of the nonlinear tuning surface qualitatively impacted a neuron’s behavior.

To quantify both the symmetry and the vertical orientation of the marginal tuning curves, we defined a tuning symmetry index (TSI), computed from the average slope and curvature of each tuning curve (Eq. 10). A TSI of 1 or -1 indicates a completely symmetric upward- or downward-facing tuning curve, respectively, and a value of 0 indicates a completely asymmetric tuning curve with a uniformly positive or negative slope. Tuning curves were grouped into terciles by their symmetry index, corresponding to peaks that occurred naturally at -1, 0, and 1 in the distribution of values (Fig. 6A, S10A-B, TSI<-0.3, - 0.3<TSI<0.3, and TSI>0.3). We then measured how the proportion of each TSI group varied with spike width and cortical depth (Fig. 6B). For regular spiking neurons, TSI was most often negative, and it did not vary substantially with depth. For narrow spiking neurons, on the other hand, TSI was more likely to be positive, especially at depths just below the layer 3-4 boundary.

**Figure 6.**
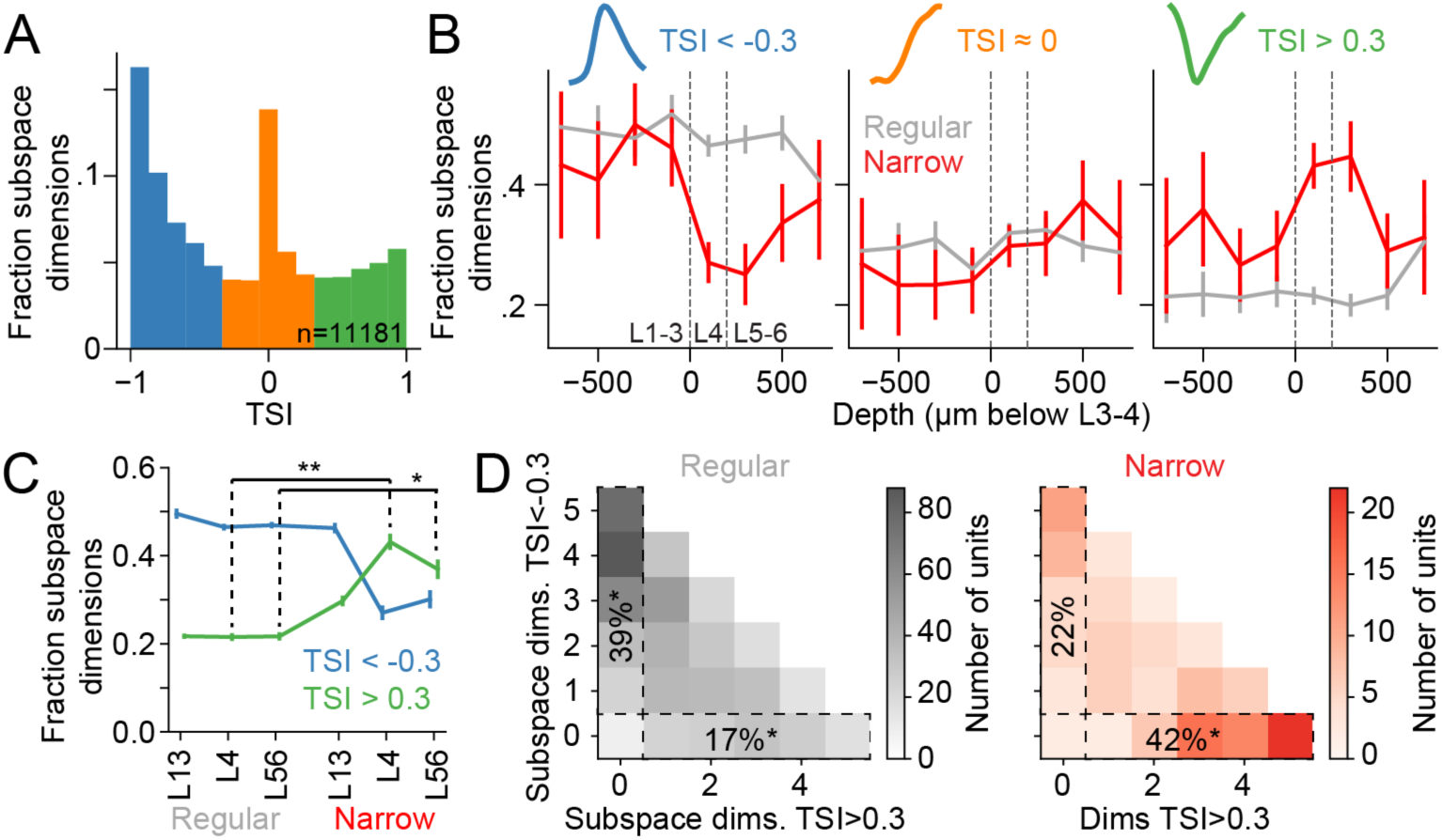
Diversity of subspace tuning nonlinearities across cell types. **A.** Distribution of TSI values measured from the marginal tuning nonlinearity of all subspace dimensions (n=1632 regular, 374 narrow AC units, 11181 unit/subspace dimension combinations, accounting for 95% of dSTRF variance for each unit). Shading indicates classifications as symmetric downward (TSI<−0.3, left), asymmetric (−0.3 < TSI < 0.3, middle), and symmetric upward (TSI>0.3). **B.** Fraction of tuning nonlinearities in each TSI category, grouped by cortical depth (x-axis) and spike width (color), for the same set as in A. Examples of nonlinearities from each category are inset above the plots. Upward symmetric nonlinearities are less common overall but more common for narrow-spiking units in cortical layer 4 (between dashed lines, generalized linear mixed effects model: spike width, χ^2^=62.0, p=3.4e-15; depth, χ^2^=3.9, p=0.14, spike width x layer, χ^2^=13.6, p=0.0011, full details in Table S2). **C.** Fraction of positive (>0.3. blue) and negative (<-0.3, green) TSI, for subspace dimensions grouped by spike width and cortical layer (see Fig. S10 for analysis including dimensions with asymmetric TSI near 0). Fraction of positive TSI is significantly greater for narrow than regular spiking neurons in layer 4 (L4, **p=7.2e-6, Z test, generalized regression model) and layers 5-6 (L56, *p=0.024). Error bars indicate 1 SEM, computed by jackknifing. **D.** Heatmaps show two-dimensional histograms of AC units binned according to their number of symmetric upward (TSI > 0.3, x-axis) and symmetric downward (TSI < -0.3, y-axis) subspace dimensions. Data for regular-spiking units at left (gray); narrow-spiking at right (red). Numbers indicate the percent of units falling in dashed boxes, with exclusively upward- or downward symmetric nonlinearities (*significantly different from chance, p<0.0001, permutation test). Data are shown only for the first five subspace dimensions (n=739 regular, 144 narrow-spiking, subset of units with at least 5 significant dimensions).

We used a generalized linear-mixed effect model to quantify the relationship between cell type (spike width and cortical depth) and the sign of TSI for each subspace dimension of each neuron (Eq. 12, detailed in Table S2). To account for the apparent nonmonotonic relationship between depth and TSI, depth was cast as a categorical variable, corresponding to layers 1-3 (depth < 0), layer 4 (0 < depth < 200) and layers 5-6 (depth > 200). The occurrence of asymmetric TSI (−0.3<TSI<0.3) was relatively constant across cortical depths. Thus, we excluded data in this category to permit training of a binary classifier for positive versus negative TSI (Fig. 6B-C, *n*=5644/8179 A1 unit/filter pairs with TSI < -0.3 or >0.3). As suggested by the data, spike width had a large regression weight (χ^2^=62.0, p=3.4e-15). There was no effect of cortical depth alone (χ^2^=3.89, p=0.14), but there was an interaction between depth and spike width, reflected primarily in layer 4 (depth 0-200, χ^2^=13.6, p=0.0011). Similar effects were observed in PEG (Table S2).

In several example units, tuning nonlinearities for the highest variance subspace dimension were more likely to be asymmetric than other dimensions (Dim. 1, Fig. 3C). TSI in this asymmetric range is consistent with the monotonic nonlinearity implemented in LN models ^45^. The tendency for the first dimension to be asymmetric was similar across all cell types (Fig. S10C-D). While the fraction of dimensions with TSI near 0 differed between first versus later subspace dimensions, the relatively greater likelihood of positive TSI for narrow spiking neurons was consistent when only dimensions with TSI < - 0.3 or > 0.3 were considered (Fig. S10E-F).

We also observed that, within a single unit, symmetric tuning nonlinearities tended to all be upward or downward facing across subspace dimensions (compare columns 2 and 3, Fig. 3C). To quantify this effect, we counted the number of subspace dimensions for each A1 unit with TSI < -0.3 or TSI > 0.3. The histogram of counts showed that units were much more likely than chance to have nonlinearities only in one of these categories (Fig. 6D, dashed boxes, permutation test, shuffling TSI values across neurons). Consistent with their tendency toward positive TSI (Fig. 6C), all the subspace dimensions for many narrow spiking neurons had only TSI > 0.3. Thus, when the subspace model revealed symmetric nonlinear tuning, the direction of the tuning curve within a single unit (upward or downward) was consistent across subspace dimensions.

## Discussion

We find that a low-dimensional sensory tuning subspace can be readily extracted from a CNN-based encoding model for neurons in auditory cortex (AC). A new encoding model, based only on the projection of stimuli into the subspace, performs nearly as well as the original CNN. Thus, this simpler and more interpretable model accounts for the key computations performed by the CNN. Analytically, the subspace model is closely related to multifilter models estimated using spike-triggered covariance (STC) and maximally informative dimensions (MID), establishing a conceptual link between CNNs and these previously proposed architectures ^2,19^. While CNNs have been shown to account for sensory coding qualitatively better than other encoding models, understanding their key computations has been challenging ^5,8^. We find that subspace models clearly describe nonlinear response properties and elucidate functional differences between neuronal cell types, distinguishing functional properties between narrow- and regular-spiking neurons and between neurons at different cortical depths (Fig. 7). The current study focuses on neurons in the auditory system, but subspace analysis is a general method that can be applied to a wide range of sensory and other neural systems.

**Figure 7.**
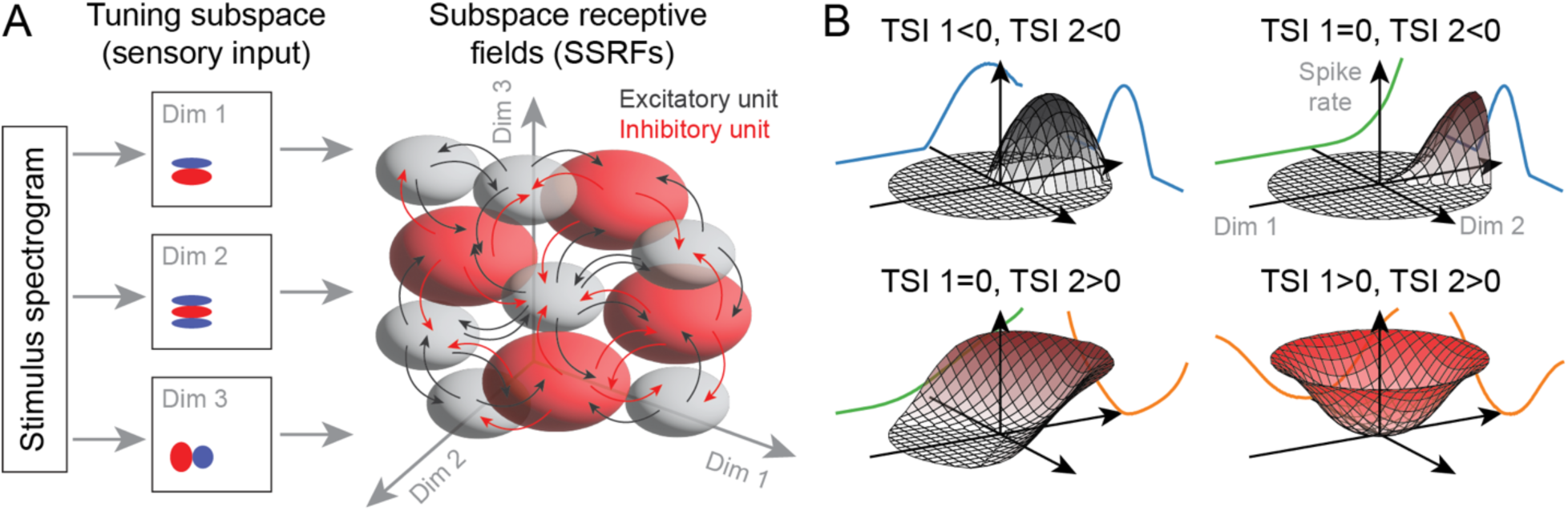
A. Conceptual framework for sparsely tiled subspace receptive fields (SSRFs) in local cortical circuits. The site-wide tuning subspace determines the range of sensory inputs reaching the local population in a cortical column. Shaded volumes indicate the maximum response areas of single units to stimuli projected into three dimensions (Dim 1-Dim 3) of the tuning subspace. While most neurons receive these inputs, strong recurrent inhibition drives each neuron to respond to distinct stimuli, reducing SSRF overlap within the shared space (Fig. 4). The larger SSRFs for inhibitory neurons (red) may reflect broadly tuned inhibition that pushes tuning of excitatory neurons (gray) to a narrower range of stimuli. **B.** Representative schematic SSRFs reflecting the observed distribution of temporal symmetry index (TSI, Fig. 6). For visualization, tuning surfaces show responses (z axis) to stimuli projected onto the first two subspace dimensions (Dim 1/Dim 2, x-/y axes). For most regular spiking neurons (top row, gray), tuning nonlinearities are mostly downward symmetric (TSI < -0.3), producing responses to a narrow range of stimuli. For some narrow-spiking neurons, nonlinearities are upward symmetric (TSI > 0.3), producing “bowl-like” tuning, with responses to many high-contrast stimuli in the tuning subspace. Neurons in both groups often have monotonic tuning along the first subspace dimension (TSI ≈ 0). The exemplar SSRFs can tile the subspace if they are shifted along Dim 1 and rotated around the z-axis.

### Selectivity within a tuning subspace forms a sparse code in local cortical populations

While it is generally agreed that neurons in the same cortical column encode similar sensory features, they can still differ substantially in their specific response properties ^46,47^. Our analysis of local neural populations in AC revealed that nearby neurons share a similar sensory tuning subspace, but their individual subspace receptive fields (SSRFs) sparsely tile that subspace (Fig. 7A). Inputs from thalamus to cortex are relatively low-dimensional and may determine the small number of dimensions in the site-wide tuning subspace. Recurrent inhibition within a much larger cortical population can drive individual neurons to respond to distinct stimuli within this subspace ^38,48^. Thus, for any single stimulus, only a small fraction of neurons will produce strong responses, while their neighbors will be suppressed. This computation is often associated with gain control mechanisms and has been proposed to produce a sparse population code ^49,50^. A sparse representation can facilitate selection of sensory features for binding features into objects and selecting information to guide behavior ^51^. This sparse tiling also accounts for the relatively low signal correlations we observe in activity evoked by natural sounds, even though nearby neurons all share basic tuning properties, like best frequency and tuning bandwidth ^47^.

### Flattened CNNs reveal sensory tuning subspaces

Understanding computations by which the brain extracts and encodes information from dynamic natural sounds is a long-standing problem in sensory neuroscience ^12^. The classic linear-nonlinear spectrotemporal receptive field (LN STRF) describes encoding as convolution of the sound spectrogram with a single linear spectrotemporal filter, followed by a static rectifying nonlinearity ^52,53^. Subspace encoding models have been proposed as a generalization of the LN STRF, in which the stimulus is convolved with two or more filters, and the response is then a nonlinear combination of the projection into this tuning subspace ^2,18,19^. The subspace extracted from a CNN describes multiple sensory features that influence neural activity and the nonlinear interactions that shape responses. Peak activations in the SSRF provide a means of determining optimal stimuli, an approach that is used to study neural DNNs ^8,54^. In addition, equal-amplitude contours on the tuning surface predict stimuli that produce an invariant response, an approach that can be tested with closed-loop experiments.

While informative, subspace models can be difficult to fit directly to neural data, especially for natural stimuli ^10^, and their implementation has been limited to 2 or 3 subspace filters ^2,25,55^. In contrast, large CNN models can be fit readily to large natural stimulus datasets ^8,11,56^. The flattening process developed in this study effectively uses large amounts of activity simulated by the CNN model to fit a subspace model. The resulting subspace model can account for time-varying neural activity nearly as accurately as the full CNN. In addition to facilitating subspace analysis, this relationship establishes a direct conceptual link between deep learning-based encoding models and multifilter encoding models.

### Tuning within sensory subspaces accounts for gain control and sensory invariance

The shape of tuning nonlinearities imposes qualitative differences in how a subspace dimension influences neural response properties ^2^. Asymmetric nonlinearities produce a response characteristic of the LN model, where spike rate changes monotonically with stimulus contrast. If the nonlinearity is symmetric, a stimulus will have the same effect on spiking, whether it projects to a large positive or negative value in the tuning subspace (Fig. 7B). In most neurons, symmetric nonlinearities were downward facing (inverted-U), meaning that large positive or negative stimulus projections produced a decrease in firing. Downward-facing nonlinearities may explain contrast gain control effects often observed in sensory cortex ^57,58^. Models implementing multiplicative context fields, have shown that gain control is spectrotemporally tuned for individual neurons ^59^. Here, the subspace model identifies a set of subspace dimensions with an inverted-U nonlinearity, suggesting that suppressive effects may be tuned to a low-dimensional subspace instead of just a single dimension. Moreover, the nonlinearities are not necessarily symmetric around 0, and thus are not simply a function of contrast. Instead, the suppressive dimensions reduce responses to stimuli that deviate from one that optimally activates the neuron, enabling local populations to tile the tuning subspace (Fig. 7).

Tuning nonlinearities in other AC neurons are symmetric and upward facing (upward-U, Fig. 7B, bottom). For these subspace dimensions, a stimulus feature projecting to a large positive or negative value produce an increase in firing. Individual dimensions are, by definition, orthogonal, but examples show that pairs of filters can lie in quadrature (Fig. 3). This organization produces responses that are tuned to a spectrotemporal feature but invariant to its phase. Complex cells in the visual cortex are described as having tuning invariant to spatial phase, but reports of analogous computations in AC have been less consistent ^60^. We observe phase invariance for both spectral and temporal modulations. In the case of modulations on the temporal axis, this computation can produce rate coding of temporally modulated sound ^61^. Phase invariance on the spectral axis (Fig. 1) produces a response to narrowband tones invariant to their precise frequency but suppressed by broadband stimuli. This form of invariance has not previously been reported for AC neurons, suggesting a new, nonlinear computation by the auditory system.

### Cell type dependence of subspace properties

In the current study, all neurons from the same AC recording site had similar subspace tuning, consistent with previous observations of topographic maps for sound features ^47,62^. However, the degree of similarity within a site depended on neuronal cell type. It is hypothesized that functional differences between cell types provide insight into computations performed by local cortical circuits ^13^. Previous work has shown cell-type dependence of frequency tuning and adaptation properties ^41,63–65^. However, the relationship between local circuit position and other spectrotemporal encoding properties is less regular, suggesting that variability is nearly as large within-as between local populations ^46,66^.

We found that the similarity of tuning subspaces was especially high between pairs of narrow spiking (putative inhibitory) neurons in superficial layers, and it was weakest for regular spiking (putative excitatory) neurons in deeper layers. Superficial inhibitory neurons receive intracortical inputs and are hypothesized to mediate top-down control of sensory processing ^67,68^. In a specific topographic location, a top-down signal may thus specifically amplify or suppress representation of stimuli in the shared tuning subspace of these superficial neurons. More diverse tuning in deeper layers may describe the range of features for which responses are also modulated by the selective top-down signal. Such a mechanism might mediate selective attention to auditory features. For example, attention to the pitch of a particular voice may amplify responses to other sound features that occur coherently with that pitch ^69^.

Nonlinear properties of the SSRF also vary with neuronal cell type (Fig. 7B). Most tuning nonlinearities were downward facing, indicating a suppression of responses consistent with contrast gain control. However, neurons with narrow spike width and positioned below the layer 3/4 boundary were much more likely to have upward facing nonlinearities. The most prominent inhibitory cell type in AC layer 4 is the parvalbumin positive (PV) interneuron ^14^. PV neurons provide feedforward inhibition of thalamic inputs that can shape the frequency tuning of nearby pyramidal cells ^13,41^. Invariance to the phase of spectrotemporal modulation may produce a broadly tuned response, leading to inhibitory gain control in their post-synaptic targets. Thus, the comprehensive characterization provided by the subspace encoding model enables linking distinct computations to specific elements of local cortical circuits. Experiments that characterize subspace tuning of neuron populations with defined connectivity, like superficial inhibitory neurons that receive top-down cortical feedback ^70^ or subcortical-projecting pyramidal neurons ^65^, will provide greater insight into the role of local circuits in shaping cortical representations. Moreover, studying if and how top-down feedback affects the size and overlap of SSRFs will provide insight into how cortical circuits shape sound representations to guide sensory behavior.

## Material and Methods

### Surgical procedures

All procedures were performed in accordance with the Oregon Health & Science University Institutional Animal Care and Use Committee (IACUC) and conform to the standards of the Association for Assessment and Accreditation of Laboratory Animal Care (AAALAC) and the United States Department of Agriculture (USDA). Four neutered young adult ferrets (3 male, 1 female) were obtained from a supplier (Marshall Farms). In each animal, sterile head-post implantation surgeries were performed under anesthesia to expose the skull over the auditory cortex (AC) and permit head-fixation during neurophysiology recordings. Surgeries were performed as previously described ^71^. After removing tissue and cleaning the skull, two stainless steel head posts were anchored along the midline using light-cured bone cement (Charisma, Kulzer). To improve implant stability, 8-10 stainless self-tapping set screws were mounted in the skull. Layers of bone cement were used to build the implant to a final shape amenable to neurophysiology and wound margin care, which included frequent cleaning and sterile bandaging. Following a two-week recovery period, animals were habituated to head-fixation and auditory stimulation.

### Acoustic stimuli

Stimuli for model fitting consisted of large sequences of natural sound segments. Segments were drawn from one of two large corpora: Audioset or Pro Sound Effects (PSE) (Core 3 Complete). The Audioset corpus is comprised of recordings from over 2 million YouTube videos ^72^. The PSE corpus is comprised of 371,089 high-quality recordings and sound effects (over 3,000 hours of audio). We selected 484,375 segments from each corpus (250k 50-ms segments, 125k 110-ms segments, 62.6k 190-ms segments, etc.). Five segment durations were used (50, 110, 190, 420, and 780 ms). The pseudo-logarithmic distribution of durations avoided exact integer multiples to minimize the likelihood of harmonic ringing in the neural responses. Each sequence was 17.79 seconds in duration and included 124 segments (64 50-ms segments, 32 110-ms segments, 16 190-ms segments, etc.) plus 5 segments of silence, one per duration. The ordering of segments was random and the sound level of segments was sampled from a uniform distribution with a 20 dB range. Segments were combined using crossfading at the boundary to avoid click artifacts (10 ms Hanning window).

Segments were selected by randomly sampling audio files without replacement from the Audioset and PSE corpus. To avoid selecting highly similar segments, an auditory excitation pattern was computed for each segment, and segments were selected to have distinct excitation patterns. The excitation pattern was computed by multiplying the power spectrum (|𝐹𝐹𝑇(𝑥)|^!^) of the segment with cosine filters designed to coarsely mimic cochlear filtering in the ferret, followed by compression (raising the outputs to the 0.3 power). The structure of the cosine filters has been described previously for humans ^73^. The only difference is that we used an alternate formula for calculating *ERN_N_* derived based on ^74^:

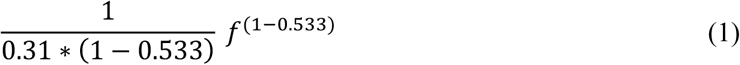

We excluded segments whose excitation pattern was similar to a previously selected segment from the same recording (r > 0.8) or whose excitation pattern was similar (r > 0.8) to more than 5% of previously selected segments (computed by randomly sampling up 10,000 previously selected segments). Audioset contains a large amount of speech and music. To prevent these stimuli from dominating the stimulus set, we prevented any one of the category labels from the Audioset ontology from comprising more than 20% of the selected Audioset segments.

For computational speed, we only excerpted segments from the first 10 seconds of each sound file (some PSE files are much longer than this duration). Stereo sounds were converted to mono by selecting the left channel, and all stimuli were resampled to 44100 Hz. For each file, we divided the recording into frames of the desired segment duration (hop equal to half the segment duration). Segment durations were 20 ms longer than those noted above to allow for crossfading. We computed the power of each frame and discarded any frames whose power was 25 dB below the maximum power across all frames in that recording (to discard silent periods). We also discarded frames that violated the excitation pattern similarity criteria described above. We then selected the segment with the greatest power from those remaining.

Stimuli for model testing consisted in part of 4 sequences created using the same approach described above. In addition, the test set contained 2, 18-second sequences composed of 1-second segments of natural and synthetic sounds tested in prior experiments from our lab ^10^. These included 6 synthetic segments comprised of tones and noises and 18 natural sound segments that included ferret vocalizations, marmoset calls, speech, music, and environmental sounds. Each sequence was composed of 12 segments with a 500-ms buffer period between each segment. Thus, in total, the test set consisted of 6 sequences with a total duration of ∼100 seconds (4*17.79+2*18=107.16). These sequences were presented between 10 and 20 times depending on the amount of recording time available.

Digital acoustic signals were transformed to analog (National Instruments), amplified (Crown), and delivered through a free-field speaker (Manger) placed 80 cm from the animal’s head, 0 ° elevation, and 30 ° contralateral to the recording hemisphere. Stimulation was controlled using custom MATLAB software (https://bitbucket.org/lbhb/baphy) and all experiments took place inside a custom double-walled sound-isolating chamber (Professional Model, Gretch-Ken). Each stimulus sequence was presented at 65 dB SPL (RMS power).

### Neurophysiology

Neurophysiological recordings were performed in awake, passively listening animals while head-fixed. To prepare for neurophysiological recordings, a small craniotomy (0.5-1mm) was opened over AC.

Recording sites were targeted based on tonotopic maps and skull landmarks ^75,76^ identified during implantation surgery. Initially, tungsten microelectrodes (FHC, 1-5MΩ) were inserted into the craniotomy to characterize tuning and response latency. Short latency responses and tonotopically organized frequency selectivity across multiple penetrations defined the location of primary auditory cortex (A1) ^75^, whereas secondary auditory cortex (posterior ectosylvian gyrus, PEG) was identified as the field ventrolateral to A1. The border between A1 and PEG was identified from the low-frequency reversal of the tonotopic gradient.

Once a cortical map was established, subsequent single-unit recordings were performed using two different electrode configurations. Experiments in animal 1 recorded neural activity using acute insertion of 64-channel silicon electrode arrays, which spanned 1.05 mm of cortical depth ^26^. Experiments in animals 2-4 recorded activity using Neuropixels short NHP probes (IMEC, 384/960 selectable channels, ^77^). Probe tips were sharpened in lab or purchased pre-sharpened to permit penetration of dura. Neuropixels probes were implanted semi-chronically using a manual microdrive (modified from ^78^) secured on the animal’s implant with dental cement. The array was implanted and allowed to settle for at least 18 hours to permit stable recordings. After recordings were complete at one site, the probe was explanted, sterilized, and reimplanted in a new craniotomy. Typically, about 150 of the 384 active channels spanned the depth of AC, as determined by current source density analysis (see below). Electrophysiological signals were amplified (RHD 128-channel headstage, Intan Technologies, or Neuropixels headstage, IMEC), digitized at 30 kHz (Open Ephys) ^79^, and saved to disk for further analysis.

Spikes were sorted offline using Kilosort2 (https://github.com/MouseLand/Kilosort2) ^80^, and sorting results were manually curated in phy (https://github.com/cortex-lab/phy). A contamination percentage was computed by measuring the cluster isolation for each sorted and curated spike cluster, which was classified as a single unit if contamination percentage was less than or equal to 5%. Clusters with contamination above 5% were classified as multi-unit and excluded from analysis.

### Laminar depth analysis

We used current source density analysis to classify units by cortical layer (1/3, supragranular; 4, granular; 5/6 infragranular). The local field potential (LFP) signal was generated by lowpass filtering either the raw signal from the 64-channel silicon probe or the LFP signal from the Neuropixel probe below 250Hz using a zero-phase shift (filter-filter method) 4^th^ order Butterworth filter, followed by down-sampling to 500Hz. A custom graphical interface was used to mark boundaries between layers, based on features of average sound-evoked LFP traces sorted by electrode depth (https://github.com/LBHB/laminar_tools). Layer-specific features included the pattern of current source density (CSD) sinks and sources evoked by best frequency-centered broadband noise bursts. Patterns were selected to match auditory evoked CSD patterns seen in AC of multiple species ^81,82,43^. Each unit was assigned a layer based on the boundaries above and below the channel where its spike had the largest amplitude.

### Spike width classification

We classified neurons as narrow- and broad-spiking based on the average width of the waveform. Width was calculated as the time between the depolarization trough and the hyperpolization peak ^39^. The distribution of spike width across neurons was bimodal, and the categorization threshold was defined as the minimum between the bimodal peaks. Filtering properties differed between 64-channel probes and Neuropixels, thus categorization threshold was defined as 0.35 ms and 0.375 ms, respectively. Consistent with studies in other systems, recordings from optogenetically labeled GABAergic neurons in ferret auditory cortex have demonstrated that these inhibitory neurons fall entirely in the narrow-spiking group^40^.

### Convolutional neural network (CNN) encoding model

Encoding model analysis was used to describe the transformation from stimulus spectrogram to neural response. The sound-evoked response was defined as the peristimulus time histogram (PSTH), *r*(*t*), a neuron’s time-varying spike rate, sampled at 100 Hz (10 ms bins). Model input was the spectrogram, *s*(*f*,*t*), of the sound waveform, computed using a second-order gammatone filter bank ^83^. The spectrogram was fixed for each model rather than fitting the spectrogram’s parameters, since we have observed little benefit on model performance from this additional complexity. The filter bank consisted of *F* = 32 filters with *f_j_* spaced logarithmically from *f*_low_ = 200 to *f*_high_ = 20,000 Hz (approximately 1/6 octave per bin).

Filter bank output was smoothed, downsampled to 100 Hz to match the sampling of the neural PSTH, and log-compressed to account for the action of the cochlea. For the stimulus spectrogram, defined as a function of sound frequency and time, 𝑠(𝑓, 𝑡), a causal encoding model prediction at time, *t*, for neuron, *i*, is then a function of the stimulus up until that time, 𝑟_+_(𝑡) = 𝐻_+_[𝑆(𝑡)], where 𝑆(𝑡) = [𝑠(𝑓, 1) . . . 𝑠(𝑓, 𝑡)].

#### Linear-nonlinear model

The linear-nonlinear spectrotemporal receptive field (LN) model is widely used in studies of neural auditory coding ^1,84,85^ and was used as a baseline reference for this study (Fig. 1). To leverage statistical power in the neural population data and permit direct comparison with the population CNN model, below, the LN model was implemented using a shared filter space across neurons in each recording site. The population LN model is analytically identical to the traditional single-neuron LN model but performs with slightly greater accuracy for neurons with low response reliability ^10^. The first stage of the population LN model convolves a bank of *M* finite impulse response (FIR) filters, *h*, with the stimulus spectrogram:

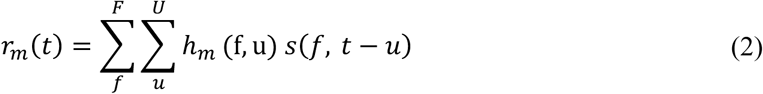

Here, we used *M*=100 filters, each containing *F* = 32 spectral channels and *U* = 25 temporal bins. Filters are implemented with rank-1 factorization. Higher rank filters could be used, but we have found that in a trade-off between larger filter bank (greater *M*) and higher-rank filters, the additional filters provide better model performance ^44^. Activity in this filtered space is then linearly weighted to generate a linear prediction for each neuron *i*:

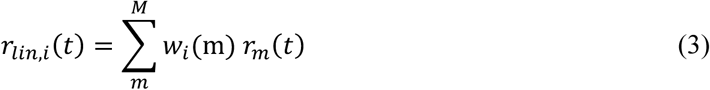

Finally, this linear prediction is passed through a double exponential nonlinearity to account for spiking threshold and firing rate saturation to produce the final prediction of the PSTH response. The double exponential nonlinearity is:

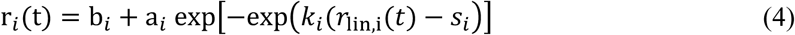

where the baseline spike rate, saturated firing rate, firing threshold, and gain are represented by *b*, *a*, *s* and *k*, respectively ^44^.

#### Population CNN model

The CNN model employed an architecture developed in a previous study of natural sound encoding in AC ^10^. Like the LN model, this architecture fit a shared subspace across neurons recorded simultaneously from the same local population. The time-varying activity of individual neurons was computed in the final layer, as the weighted sum of outputs from the penultimate (shared) model layer ^8,10^. The full model consisted of two convolutional blocks, followed by two dense layers.

Each convolutional block was composed of a dense layer and a set of finite-impulse-response (FIR) filters with ReLU activation. The FIR filters operated on each output of the dense layer separately. Thus, each convolutional block effectively performed the computation of a spectrotemporal receptive field (STRF) in the standard LN model ^10^. The first block contained 80 filters that each integrated across the 32 spectrogram channels and then performed convolution over the 15 preceding 10-ms time bins (80 32x15 filters, Eq. 2). The second block contained 100 80x10 linear filters. These convolutional blocks were followed by two densely connected layers: 1) 100x100 with ReLU activation, and 2) 100x*N* with a double exponential nonlinearity, identical to that used in the LN model (Eq. 4) and fit individually for each neuron (N: number of neurons being fit).

### Model optimization

All fitting was performed exclusively with the estimation dataset, and the validation set was used only for measuring prediction accuracy. Prior to fitting, all input (stimulus) and output (time-varying spike rate) channels were normalized to a range of 0 to 1. Model parameters were then fit using TensorFlow’s implementation of the Adam algorithm for stochastic gradient descent ^86^, using a mean squared error (MSE) loss function, with L2 regularization (alpha=10^-4^). Loss was computed for all neurons in the dataset simultaneously. We evaluated the use of a Poisson loss function for model optimization ^30^ and observed similar but slightly lower prediction accuracy than for models trained with MSE loss. To obtain an initial coarse fit and prevent falling into local minima, the final double exponential nonlinearity was replaced with a simple level shift, and the fit was performed with a relatively high learning rate (0.01) and stop tolerance for fit loss (0.001). The double exponential was then restored and fitting continued with smaller learning rate (0.001) and tolerance for fit loss (10^-4^). To mitigate overfitting, an 8-fold jackknifing procedure was used, in which a distinct 12.5% of the fit set was excluded from the fit data set for each model fit. For each jackknife sample, the model was fit starting at 5 random initial conditions, making 40 model fits in total. The highest performing model for each jackknife was saved for subsequent analysis.

### Tuning subspace analysis

#### Dynamic spectrotemporal receptive field (dSTRF) measurement

While the population CNN model describes a complex nonlinear function, the stimulus-response relationship at each time point during stimulation can be approximated as a linear spectrotemporal receptive field, termed the dSTRF. This locally linear approximation can be measured as the derivative of the response predicted by the CNN model for the *i*-th neuron at each timepoint, *t*, r_+_(𝑡) = H_+_[𝑆(𝑓, 0 … 𝑡)], relative to the input stimulus spectrogram, 𝑆(𝑓, 𝑡),

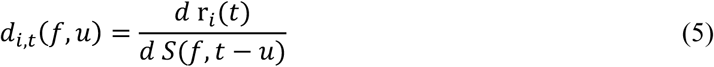

Intuitively, the dSTRF is a locally linear approximation of a nonlinear function, analogous to fitting an LN model using only small perturbations in the stimulus at each frequency, *f*, and time lag, *u*. The dSTRF was measured with a maximum time lag, *U*=250 ms, which corresponds to the maximum integration time of the population CNN. We measured the dSTRF by computing the Jacobian of the population CNN output at each fit stimulus timepoint, which is equivalent to Eq. 5. To reduce noise, the dSTRF was measured using each of the 8 jackknife model estimates. A shrinkage function (soft threshold for mean/standard error<1) was used to attenuate noise in the dSTRF average across jackknifes ^87^. Thus, for a single neuron and stimulus of length (in samples) *T*, the final dSTRF consisted of *T* linear spectrotemporal filters, 𝑑_+,8_(𝑓, 𝑢).

#### Subspace encoding model

This model was derived from the CNN to determine if the CNN captured the same functional properties as previously proposed subspace models ^2,30,32^. The large collection of dSTRFs measured for each neuron was projected into a low dimensional space by principal component analysis (PCA). The frequency, *f*, and time lag, *u*, dimensions of the dSTRF were reshaped into a single axis, and PCA was performed along the complementary time axis, *t*. A low-dimensional subspace was then defined by the *N* components explaining the most variance in the dSTRF, 𝑔_+,9_(𝑓, 𝑢), 𝑗 = 1. . . 𝑁. For most analysis, we chose *N* to account for 95% of variance, but different threshold values were assessed in some analyses. For subsequent analysis, the stimulus was projected into this low dimensional subspace,

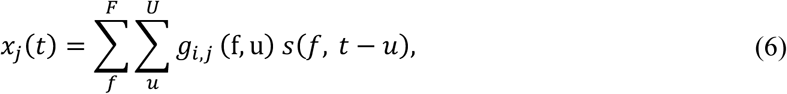

Nonlinear tuning in the subspace was visualized as a subspace receptive field (SSRF), the mean actual or predicted firing rate at each point in that space. To determine how well the tuning subspace accounted for sensory responses, the stimulus was projected into the subspace, and a densely connected three-layer neural network was used to model the relationship between the time-varying stimulus projection and neural activity in the fit data. Each layer contained 15 linear units, followed by a nonlinearity (layers 1-2: rectified linear unit; layer 3: difference of exponentials sigmoid). Thus the network could fit an arbitrary SSRF,

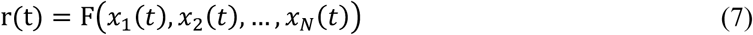

This model could account for many nonlinear interactions within the subspace but could not account for any spectrotemporal tuning outside of the *N* subspace dimensions. We tested several variants of the model, including additional dense layers and more units per layer, but no improvement in prediction accuracy was observed for larger architectures.

#### Second-order subspace model

Multifilter models using spike-triggered covariance use a second order model to define the relationship between a subspace representation and the neural response. To implement this architecture, we used a second order model to fit the mapping between stimulus subspace and time-varying response. For the *N*-dimensional subspace stimulus, *x*_2_(*t*) ... *x_N_*(*t*), the second order model response is:

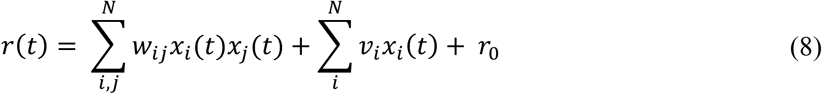

Second-order coefficients, 𝑤_+9_, are weights applied to products of values in the subspace channels *i* and *j*. When *i*=*j*, a positive value indicates an increase in response when the projection on that dimension has either a large positive or negative value, and a negative value indicates a decreased response.

#### Subspace similarity index (SSI)

To measure the similarity of tuning between two sets of *N* subspace filters, *i* and *j*, we computed the sum of the cosine similarity (CS) between each pair of filters and normalized by the number of filters.

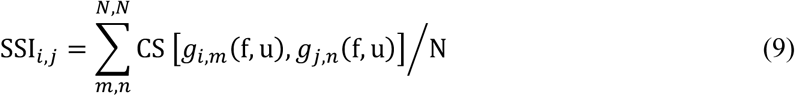

The summed cosine similarity across dimensions is an implementation of Grassman distances used for measuring the similarity of subspace projections ^88^. For the current study, we measured SSI between the *N*=4 largest filters from each neuron.

#### Tuning symmetry index (TSI)

To characterize the shape of the marginal tuning curve for a single subspace dimension, TSI distinguished between symmetric downward facing (values near -1), symmetric upward facing (values near 1) and asymmetric (values near 0). For tuning curve, 𝑦(𝑥), with derivative, 𝑦:(𝑥), and second derivative, 𝑦^::^(𝑥), we define the TSI,

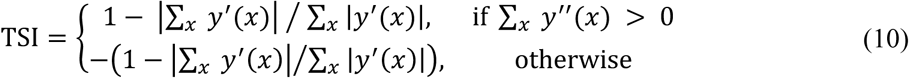

In cases where 𝑦′ is always positive or negative (asymmetric), the ratio of sums reduces to 1 and thus gives a TSI value of 0. If 𝑦′ has equal amounts positive and negative (symmetric), the ratio will be zero, and the average curvature, captured by 𝑦^::^(𝑥), will determine if TSI is -1 or +1. The derivative was approximated by discretizing the tuning curve with 20 bins spanning 99% of the values in x spanned by the stimulus projection onto that dimension.

### Statistical analysis

For all pairwise statistical tests we performed a Wilcoxon signed-rank test (sign test), and for unpaired comparisons we used a Mann-Whitney U test. Significance was determined at the alpha = 0.05 level. Units were considered sound-responsive if either the LN or CNN model predicted test responses above chance (p<0.05, permutation test, corrected for multiple comparisons, n=2337/2874 units).

Linear mixed-effects models (lme4 package in R version 4.4.3) were used to assess relationships between neuronal cell type and SSI and TSI metrics. For the SSI model (fit using the lme4::*lmer function*), the outcome was SSI for a pair of neurons, *i* and *j* (Table S1). Inputs were mean cortical depth, *D* (−750-750 μm relative to the layer 3-4 boundary), depth difference between neurons, Δ, and spike width category W (narrow-narrow, narrow-regular, regular-regular), and interactions between depth and width variables (*β_DW_*, *β_ΔW_*):

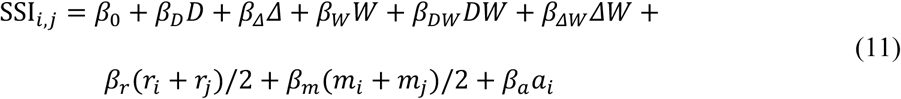

CNN prediction accuracy (*r*) and mean rate (*m*) were also included as predictors, and animal identity was included as a random effect (*a*). Significance of individual parameters was determined by a *t test*, and goodness of fit was measured by a χ^2^ test (Type II Wald using the car::Anova package). Prior to statistical modeling, continuous predictors (*D*, *Δ*, *m*, *r*) were normalized (using the *scale* function in R).

For TSI, we used generalized mixed-effect models (lme4::glmer function) with a logistic regression link function (Table S2). The outcome was whether TSI was positive (TSI > 0.3) or negative (TSI < -0.3), and data with intermediate TSI values were excluded. Predictors were neuron depth (*L*, corresponding to cortical layers 1-3 (*di* < 0 μm), layer 4 (0 < *d_i_* < 200), and layers 5-6 (*d_i_* > 200) layer), spike width (*w*), mean rate, (*m*) and CNN prediction accuracy (*r*).

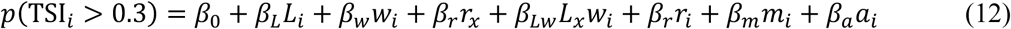

Animal identity was included as a random effect (*a*). Significance of individual parameters was determined by a *z test*, and goodness of fit was measured by a χ2 test (Type II Wald using the car::Anova package). Prior to statistical modeling, continuous predictors (mean rate and CNN prediction accuracy) were normalized (using the *scale* function in R).

## Data and code availability

All data used in this study is available open-access in Zenodo (https://zenodo.org/records/17211353).

Analysis was performed using the Neural Encoding Model System library (https://github.com/LBHB/NEMS/). Scripts to generate the study figures will be included in the data repository.

## Acknowledgements

This work was supported by NIH BRAIN Initiative Grant R01EB028155 and NIDCD Grant R01DC014950 (S.V.D.)

## References

1. Eggermont, J. J., Aertsen, A. M. & Johannesma, P. I. Prediction of the responses of auditory neurons in the midbrain of the grass frog based on the spectro-temporal receptive field. Hear. Res. 10, 191–202 (1983).

2. Atencio, C. A., Sharpee, T. O. & Schreiner, C. E. Cooperative nonlinearities in auditory cortical neurons. Neuron 58, 956–966 (2008).

3. Willmore, B. D., Schoppe, O., King, A. J., Schnupp, J. W. & Harper, N. S. Incorporating Midbrain Adaptation to Mean Sound Level Improves Models of Auditory Cortical Processing. J. Neurosci. 36, 280–9 (2016).

4. Sadagopan, S., Kar, M. & Parida, S. Quantitative models of auditory cortical processing. Hear. Res. 429, 108697 (2023).

5. Yamins, D. & Hong, H. Performance-optimized hierarchical models predict neural responses in higher visual cortex. Proc. … (2014).

6. Kell, A. J. E., Yamins, D. L. K., Shook, E. N., Norman-Haignere, S. V. & McDermott, J. H. A Task-Optimized Neural Network Replicates Human Auditory Behavior, Predicts Brain Responses, and Reveals a Cortical Processing Hierarchy. Neuron 98, 630–644.e16 (2018).

7. Giordano, B. L., Esposito, M., Valente, G. & Formisano, E. Intermediate acoustic-to-semantic representations link behavioral and neural responses to natural sounds. Nat. Neurosci. 26, 664–672 (2023).

8. Walker, E. Y. et al. Inception loops discover what excites neurons most using deep predictive models. Nat. Neurosci. 22, 2060–2065 (2019).

9. Keshishian, M. et al. Estimating and interpreting nonlinear receptive field of sensory neural responses with deep neural network models. eLife 9, e53445 (2020).

10. Pennington, J. R. & David, S. V. A convolutional neural network provides a generalizable model of natural sound coding by neural populations in auditory cortex. PLoS Comput. Biol. 19, e1011110 (2023).

11. Oliver, M., Winter, M., Dupré la Tour, T., Eickenberg, M. & Gallant, J. L. A biologically-inspired hierarchical convolutional energy model predicts V4 responses to natural videos. bioRxiv 2024–12 (2024).

12. Wu, M. C.-K., David, S. V. & Gallant, J. L. Complete functional characterization of sensory neurons by system identification. Annu. Rev. Neurosci. 29, 477–505 (2006).

13. Blackwell, J. M. & Geffen, M. N. Progress and challenges for understanding the function of cortical microcircuits in auditory processing. Nat. Commun. 8, 2165 (2017).

14. Studer, F. & Barkat, T. R. Inhibition in the auditory cortex. Neurosci. Biobehav. Rev. 132, 61–75 (2022).

15. Feather, J., Leclerc, G., Mądry, A. & McDermott, J. H. Model metamers reveal divergent invariances between biological and artificial neural networks. Nat. Neurosci. 26, 2017–2034 (2023).

16. Lundberg, S. M. & Lee, S.-I. A unified approach to interpreting model predictions. Adv. Neural Inf. Process. Syst. 30, (2017).

17. Hu, E. J. et al. Lora: Low-rank adaptation of large language models. ICLR 1, 3 (2022).

18. Brenner, N., Bialek, W. & de Ruyter van Steveninck, R. Adaptive rescaling maximizes information transmission. Neuron 26, 695–702 (2000).

19. Pillow, J. W. & Simoncelli, E. P. Dimensionality reduction in neural models: an information-theoretic generalization of spike-triggered average and covariance analysis. J. Vis. 6, 9–9 (2006).

20. Christianson, G. B., Sahani, M. & Linden, J. F. The consequences of response nonlinearities for interpretation of spectrotemporal receptive fields. J Neurosci 28, 446–455 (2008).

21. Kaardal, J. T., Theunissen, F. E. & Sharpee, T. O. A Low-Rank Method for Characterizing High-Level Neural Computations. Front. Comput. Neurosci. 11, 68 (2017).

22. Rust, N. C., Schwartz, O., Movshon, J. A. & Simoncelli, E. P. Spatiotemporal elements of macaque v1 receptive fields. Neuron 46, 945–956 (2005).

23. Kozlov, A. S. & Gentner, T. Q. Central auditory neurons have composite receptive fields. Proc. Natl. Acad. Sci. 113, 1441–1446 (2016).

24. Schinkel-Bielefeld, N., David, S. V., Shamma, S. A. & Butts, D. A. Inferring the role of inhibition in auditory processing of complex natural stimuli. J. Neurophysiol. 107, 3296–3307 (2012).

25. Hullett, P. W., Hamilton, L. S., Mesgarani, N., Schreiner, C. E. & Chang, E. F. Human Superior Temporal Gyrus Organization of Spectrotemporal Modulation Tuning Derived from Speech Stimuli. J. Neurosci. 36, 2014–26 (2016).

26. Du, J., Blanche, T. J., Harrison, R. R., Lester, H. A. & Masmanidis, S. C. Multiplexed, High Density Electrophysiology with Nanofabricated Neural Probes. PLOS ONE 6, e26204 (2011).

27. Juavinett, A. L., Bekheet, G. & Churchland, A. K. Chronically implanted Neuropixels probes enable high-yield recordings in freely moving mice. eLife 8, e47188 (2019).

28. Sabat, M. et al. Neurons in auditory cortex integrate information within constrained temporal windows that are invariant to the stimulus context and information rate. bioRxiv 2025–02 (2025).

29. Tjoa, E. & Guan, C. A survey on explainable artificial intelligence (xai): Toward medical xai. IEEE Trans. Neural Netw. Learn. Syst. 32, 4793–4813 (2020).

30. McFarland, J. M., Cui, Y. & Butts, D. A. Inferring nonlinear neuronal computation based on physiologically plausible inputs. PLoS Comput. Biol. 9, e1003143 (2013).

31. Meyer, A. F., Williamson, R. S., Linden, J. F. & Sahani, M. Models of Neuronal Stimulus-Response Functions: Elaboration, Estimation, and Evaluation. Front. Syst. Neurosci. 10, 109 (2017).

32. Real, E., Asari, H., Gollisch, T. & Meister, M. Neural circuit inference from function to structure. Curr. Biol. 27, 189–198 (2017).

33. Merzenich, M. M., Knight, P. L. & Roth, G. L. Representation of cochlea within primary auditory cortex in the cat. J. Neurophysiol. 38, 231–249 (1975).

34. Narayanan, D. P., Tsukano, H., Kline, A. M., Onodera, K. & Kato, H. K. Biological constraints on stereotaxic targeting of functionally-defined cortical areas. Cereb. Cortex 33, 3293–3310 (2023).

35. Imig, T. J. & Brugge, J. F. Sources and terminations of callosal axons related to binaural and frequency maps in primary auditory cortex of the cat. J. Comp. Neurol. 182, 637–660 (1978).

36. Schreiner, C. E. & Mendelson, J. R. Functional topography of cat primary auditory cortex: distribution of integrated excitation. J. Neurophysiol. 64, 1442–1459 (1990).

37. Schulze, H., Hess, A., Ohl, F. W. & Scheich, H. Superposition of horseshoe-like periodicity and linear tonotopic maps in auditory cortex of the Mongolian gerbil. Eur. J. Neurosci. 15, 1077–1084 (2002).

38. Wood, K. C., Blackwell, J. M. & Geffen, M. N. Cortical inhibitory interneurons control sensory processing. Curr. Opin. Neurobiol. 46, 200–207 (2017).

39. Trainito, C., von Nicolai, C., Miller, E. K. & Siegel, M. Extracellular Spike Waveform Dissociates Four Functionally Distinct Cell Classes in Primate Cortex. Curr. Biol. 29, 2973–2982.e5 (2019).

40. López Espejo, M. & David, S. V. A sparse code for natural sound context in auditory cortex. Curr. Res. Neurobiol. 6, 100118 (2024).

41. Moore, A. K. & Wehr, M. Parvalbumin-Expressing Inhibitory Interneurons in Auditory Cortex Are Well-Tuned for Frequency. J. Neurosci. 33, 13713–13723 (2013).

42. Lakatos, P., Chen, C.-M. M., O’Connell, M. N., Mills, A. & Schroeder, C. E. Neuronal oscillations and multisensory interaction in primary auditory cortex. Neuron 53, 279–292 (2007).

43. Mendoza-Halliday, D. et al. A ubiquitous spectrolaminar motif of local field potential power across the primate cortex. Nat. Neurosci. 27, 547–560 (2024).

44. Thorson, I. L., Liénard, J. & David, S. V. The essential complexity of auditory receptive fields. PLoS Comput. Biol. 11, e1004628 (2015).

45. Homma, N. Y. et al. Receptive-field nonlinearities in primary auditory cortex: a comparative perspective. Cereb. Cortex 34, bhae364 (2024).

46. Atencio, C. A. & Schreiner, C. E. Laminar diversity of dynamic sound processing in cat primary auditory cortex. J. Neurophysiol. 103, 192–205 (2010).

47. Kanold, P. O., Nelken, I. & Polley, D. B. Local versus global scales of organization in auditory cortex. Trends Neurosci. 37, 502–510 (2014).

48. Coultrip, R., Granger, R. & Lynch, G. A cortical model of winner-take-all competition via lateral inhibition. Neural Netw. 5, 47–54 (1992).

49. Olshausen, B. A. & Field, D. J. Emergence of simple-cell receptive field properties by learning a sparse code for natural images. Nature 381, 607–609 (1996).

50. Willmore, B. D. B. & King, A. J. Auditory Cortex: Representation through Sparsification? Curr. Biol. 19, R1123–R1125 (2009).

51. Vinje, W. E. & Gallant, J. L. Sparse coding and decorrelation in primary visual cortex during natural vision. Science 287, 1273–1276 (2000).

52. Theunissen, F. E., Sen, K. & Doupe, A. J. Spectral-temporal receptive fields of non-linear auditory neurons obtained using natural sounds. J. Neurosci. 20, 2315–2331 (2000).

53. Miller, L. M., Escabi, M. A., Read, H. L. & Schreiner, C. E. Spectrotemporal receptive fields in the lemniscal auditory thalamus and cortex. J Neurophysiol 87, 516–527 (2002).

54. Yamins, D. L. K. & DiCarlo, J. J. Using goal-driven deep learning models to understand sensory cortex. Nat. Neurosci. 19, 356–65 (2016).

55. Touryan, J., Felsen, G. & Dan, Y. Spatial structure of complex cell receptive fields measured with natural images. Neuron 45, 781–791 (2005).

56. Drakopoulos, F. et al. Modelling neural coding in the auditory midbrain with high resolution and accuracy. *Nat*. Mach. Intell. 1–16 (2025).

57. Barbour, D. L. & Wang, X. Contrast tuning in auditory cortex. Science 299, 1073–1075 (2003).

58. Rabinowitz, N. C., Willmore, B. D. B. B., Schnupp, J. W. H. H. & King, A. J. Contrast gain control in auditory cortex. Neuron 70, 1178–91 (2011).

59. Williamson, R. S., Ahrens, M. B., Linden, J. F. & Sahani, M. Input-Specific Gain Modulation by Local Sensory Context Shapes Cortical and Thalamic Responses to Complex Sounds. Neuron 91, 467–481 (2016).

60. Tian, B., Kuśmierek, P. & Rauschecker, J. P. Analogues of simple and complex cells in rhesus monkey auditory cortex. Proc. Natl. Acad. Sci. 110, 7892–7897 (2013).

61. Liang, L., Lu, T. & Wang, X. Neural representations of sinusoidal amplitude and frequency modulations in the primary auditory cortex of awake primates. J Neurophysiol 87, 2237–2261 (2002).

62. Schreiner, C. E. & Winer, J. A. Auditory cortex mapmaking: principles, projections, and plasticity. Neuron 56, 356–65 (2007).

63. Li, L. et al. Differential Receptive Field Properties of Parvalbumin and Somatostatin Inhibitory Neurons in Mouse Auditory Cortex. Cereb. Cortex 25, 1782–1791 (2015).

64. Natan, R. G., Rao, W. & Geffen, M. N. Cortical interneurons differentially shape frequency tuning following adaptation. Cell Rep. 21, 878–890 (2017).

65. Williamson, R. S. & Polley, D. B. Parallel pathways for sound processing and functional connectivity among layer 5 and 6 auditory corticofugal neurons. eLife 8, (2019).

66. Eggermont, J. J. Representation of spectral and temporal sound features in three cortical fields of the cat. Similarities outweigh differences. J. Neurophysiol. 80, 2743–64 (1998).

67. Pi, H.-J. et al. Cortical interneurons that specialize in disinhibitory control. Nature 503, 521–524 (2013).

68. Naumann, L. B., Hertäg, L., Müller, J., Letzkus, J. J. & Sprekeler, H. Layer-specific control of inhibition by NDNF interneurons. Proc. Natl. Acad. Sci. 122, e2408966122 (2025).

69. Shamma, S. A., Elhilali, M. & Micheyl, C. Temporal coherence and attention in auditory scene analysis. Trends Neurosci. 34, 114–23 (2011).

70. Sweeney, C. G. et al. Reliable sensory processing of superficial cortical interneurons is modulated by behavioral state. Cell Rep. 44, (2025).

71. Slee, S. J. & David, S. V. Rapid Task-Related Plasticity of Spectrotemporal Receptive Fields in the Auditory Midbrain. J. Neurosci. 35, 13090–13102 (2015).

72. Gemmeke, J. F. et al. Audio Set: An ontology and human-labeled dataset for audio events. in 2017 IEEE International Conference on Acoustics, Speech and Signal Processing (ICASSP) 776–780 (2017). doi:10.1109/ICASSP.2017.7952261.

73. McDermott, J. H. & Simoncelli, E. P. Sound texture perception via statistics of the auditory periphery: evidence from sound synthesis. Neuron 71, 926–40 (2011).

74. Sumner, C. J. & Palmer, A. R. Auditory nerve fibre responses in the ferret. Eur. J. Neurosci. 36, 2428–2439 (2012).

75. Bizley, J. K., Nodal, F. R., Nelken, I. & King, A. J. Functional organization of ferret auditory cortex. Cereb. Cortex N. Y. N 1991 15, 1637–1653 (2005).

76. Atiani, S. et al. Emergent selectivity for task-relevant stimuli in higher-order auditory cortex. Neuron 82, 486–499 (2014).

77. Jun, J. J. et al. Fully Integrated Silicon Probes for High-Density Recording of Neural Activity. Nature 551, 232–236 (2017).

78. Vöröslakos, M., Petersen, P. C., Vöröslakos, B. & Buzsáki, G. Metal microdrive and head cap system for silicon probe recovery in freely moving rodent. eLife 10, e65859 (2021).

79. Siegle, J. H. et al. Open Ephys: an open-source, plugin-based platform for multichannel electrophysiology. J. Neural Eng. 14, 045003 (2017).

80. Pachitariu, M., Steinmetz, N., Kadir, S., Carandini, M. & Harris, K. Fast and accurate spike sorting of high-channel count probes with KiloSort. in (2016).

81. Maier, A., Adams, G., Aura, C. & Leopold, D. Distinct Superficial and Deep Laminar Domains of Activity in the Visual Cortex during Rest and Stimulation. Front. Syst. Neurosci. 4, (2010).

82. Schaefer, M. K., Hechavarría, J. C. & Kössl, M. Quantification of mid and late evoked sinks in laminar current source density profiles of columns in the primary auditory cortex. Front. Neural Circuits 9, (2015).

83. Lyon, R. F. & Mead, C. An analog electronic cochlea. IEEE Trans. Acoust. Speech Signal Process. 36, 1119–1134 (1988).

84. Theunissen, F. E. et al. Estimating spatial temporal receptive fields of auditory and visual neurons from their responses to natural stimuli. Netw. Comput. Neural Syst. 12, 289–316 (2001).

85. Calabrese, A., Schumacher, J. W., Schneider, D. M., Paninski, L. & Woolley, S. M. N. A generalized linear model for estimating spectrotemporal receptive fields from responses to natural sounds. PloS One 6, e16104 (2011).

86. Abadi, M. et al. Tensorflow: a system for large-scale machine learning. in OSDI vol. 16 265–283 (2016).

87. Brillinger, D. J. Some uses of cumulants in wavelet analysis. J. Nonparametric Stat. 6, 93–114 (1996).

88. Washizawa, Y. & Hotta, S. Mahalanobis Distance on Extended Grassmann Manifolds for Variational Pattern Analysis. IEEE Trans. Neural Netw. Learn. Syst. 25, 1980–1990 (2014).

