## Supplementary material for "Convolutional neural network models describe the encoding subspace of local circuits in auditory cortex": Fig. S

**Contents:** 10 Figures, 2 Tables

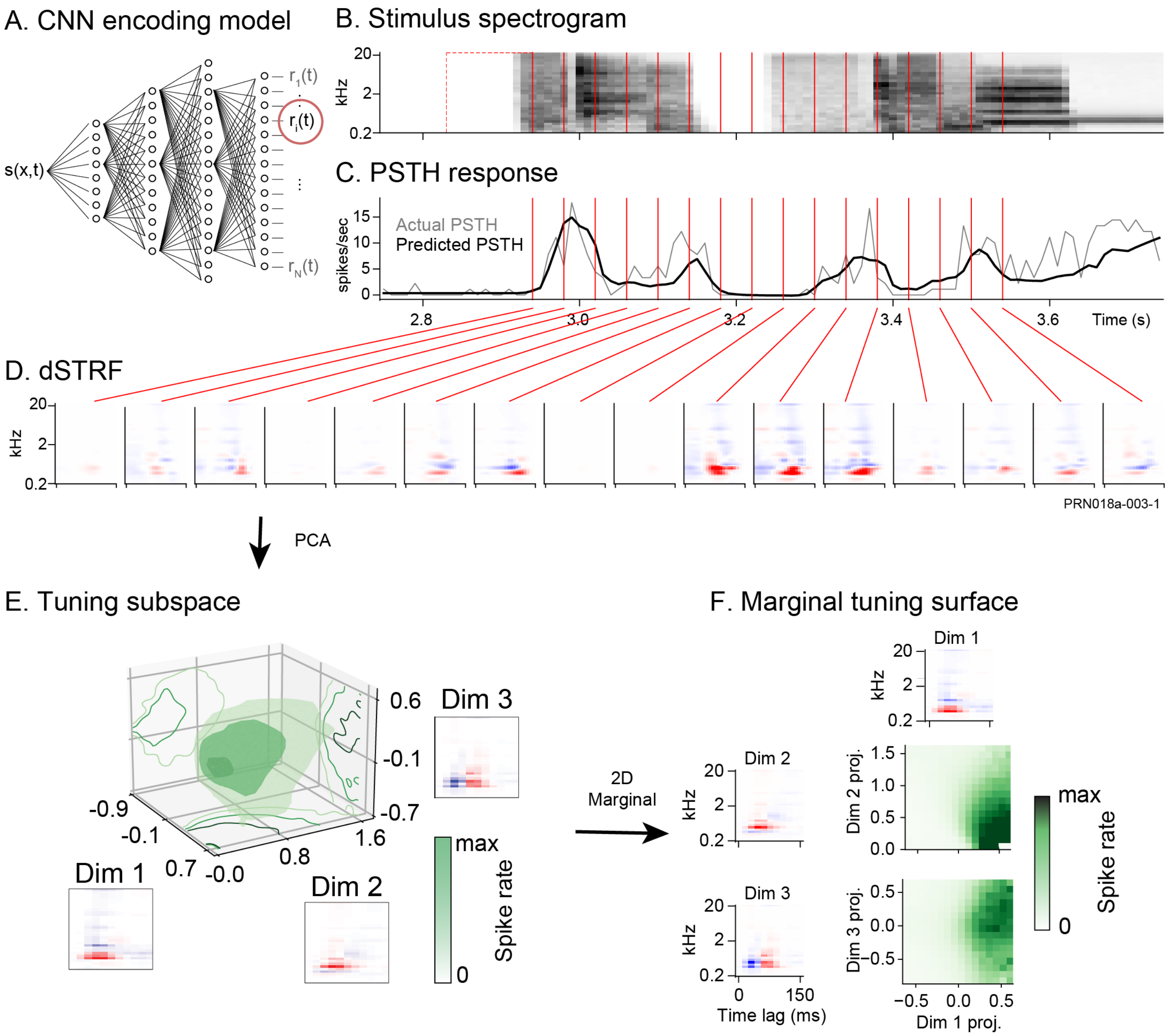


**Figure S1. A.** Natural sound encoding by single unit neurons recorded from ferret auditory cortex was modeled using a population convolutional neural network (CNN). **B.** Spectrogram of a natural sound sequence presented during one experiment. **C.** Time-varying peri-stimulus time histogram (PSTH) response recorded from one neuron during presentation of stimulus in B (gray). PSTH predicted by the CNN model overlaid in black. **D.** Dynamic spectro-temporal receptive field (dSTRF) is calculated as the derivative of the model output relative to the input stimulus. Panels show dSTRF calculated for the unit in B at example time points (vertical bars in C). **E.** PCA is applied to the large collection of dSTRFs to compute a small number of spectrotemporal filters that define the tuning subspace, i.e., that span the spectro-temporal domain of stimuli that influence neural activity. The subspace encoding model is the nonlinear function that predicts neural activity from the stimulus projection into the subspace. Red/blue heatmaps show first three PCA dimensions for the example unit. Green heatmap shows a 3D rendering of the subspace receptive field in those three dimensions. **F.** Heatmaps show response rate as a function of projection onto each of the three largest subspace dimensions (green indicates higher spike rate).

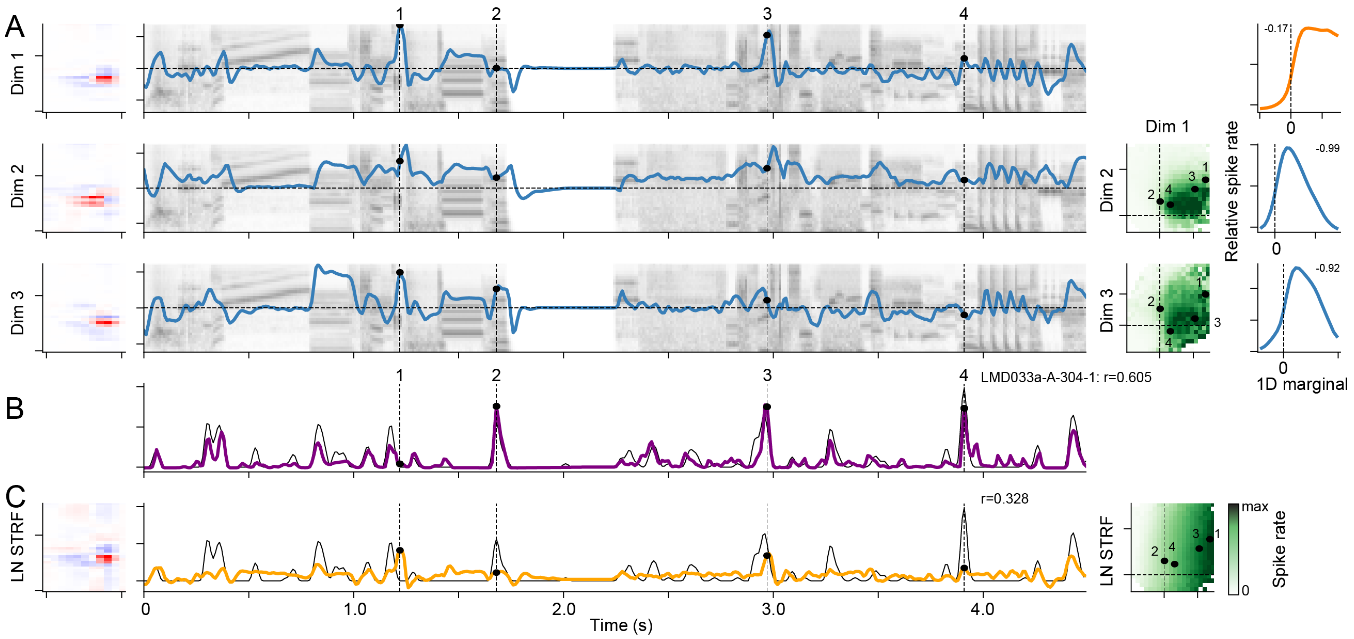


**Figure S2.** Detailed example of subspace encoding model, plotted as in Fig. 1. **A.** Subspace filters (left) are convolved with the stimulus spectrogram (second column, gray shading) to produce a projection into each dimension (3/7 significant filters shown). Tuning surface indicates the predicted response at that moment in time (third column, green shading). Numbers indicate subspace positions at example points the time-varying projections (dashed vertical lines at left). Averages across the tuning surface show mean response as a function of the projection onto each tuning dimension (fourth column). For this neuron, the first dimension has an asymmetric, monotonic nonlinearity, and dimensions 2 and 3 have a symmetric, suppressive nonlinearity. **B.** Subspace model prediction (purple) overlaid with the actual peri-stimulus time histogram (PSTH) response (gray, r=0.605). **C.** LN model fit for the same neuron (left), LN model prediction (orange) overlaid with actual PSTH (middle, r=0.328), and average LN model prediction for Dim 1 vs. Dim 2 projections in the subspace model (right).

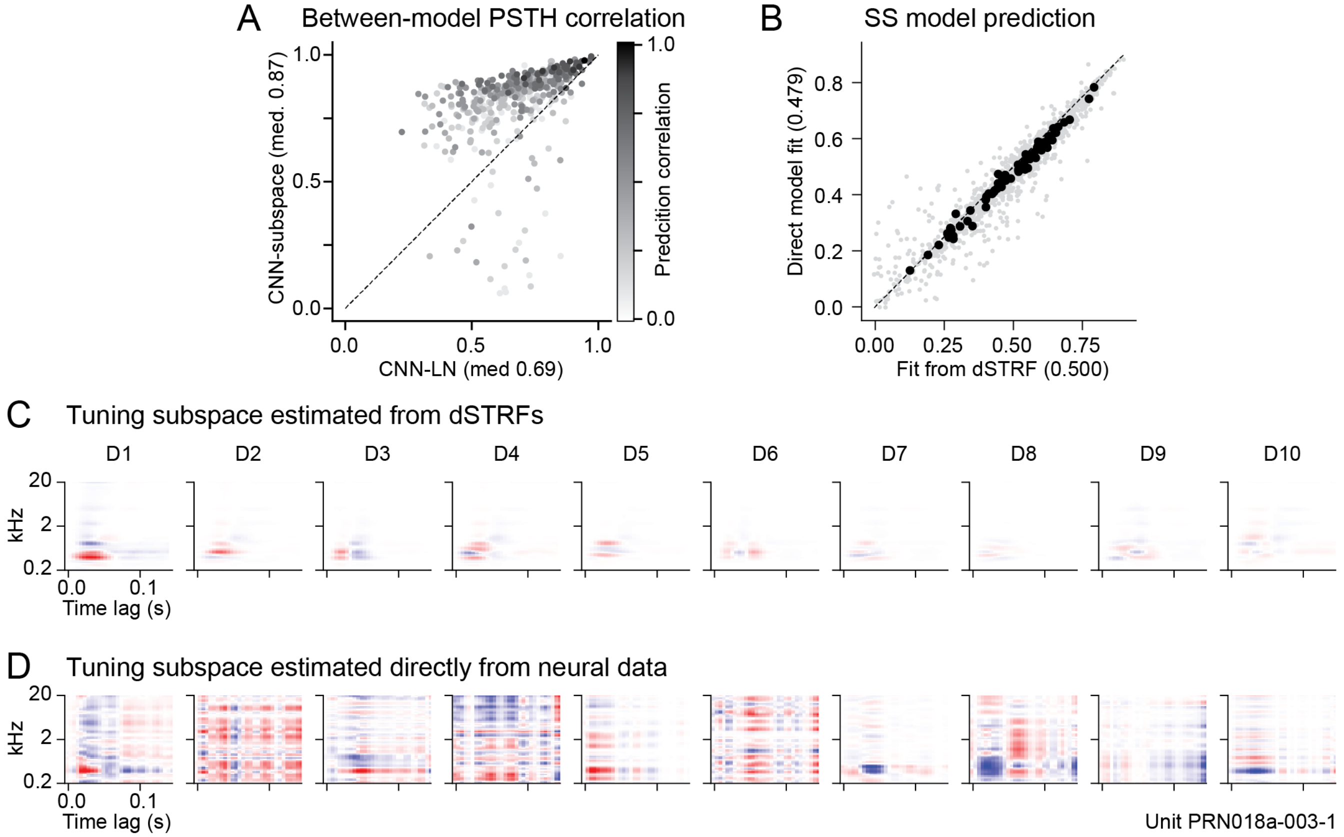


**Figure S3. A.** Functional equivalence between encoding models, computed as the correlation coefficient between their predicted PSTHs in the test dataset. Scatter plot compares correlation coefficient between predictions by CNN and LN models (x-axis, median 0.69) versus by CNN and subspace models (y-axis, median 0.87, n=1640, p=6.7e-217, Wilcoxon sign test). **B.** Scatter plot compares prediction correlation between subspace model estimated using the dSTRF (derived from the CNN, as in the main text) versus a subspace model with identical architecture (i.e., same number of subspace filters, followed by a nonlinearity) but estimated directly from the neural data. Direct training was performed with a method as closely matched as possible to the dSTRF-based model, and accuracy was measured with the same held-out natural sound dataset. While the direct fit can theoretically perform as well as the dSTRF-based estimate, accuracy is higher for the dSTRF-based model (median 0.500 versus 0.479, $n$=1267, $p$=2.1e-203, sign test). This result suggests that the dSTRF-based method regularizes the model fit more effectively than the direct fit. **C.** Ten largest subspace dimensions estimated using the dSTRF method for the example A1 neuron in Fig. 1. Heatmaps are scaled according to relative dSTRF variance explained by each dimension, providing a sense of how much each dimension accounts for evoked activity. **D.** Ten subspace dimensions from the direct-fit model for the same neuron. A broadly similar pattern of spectrotemporal tuning is observed, although filters defining the dimensions appear noisier overall for the direct fit model. Because subspace filter variance is not quantified for this fit, heatmaps were not scaled to indicate relative influence.

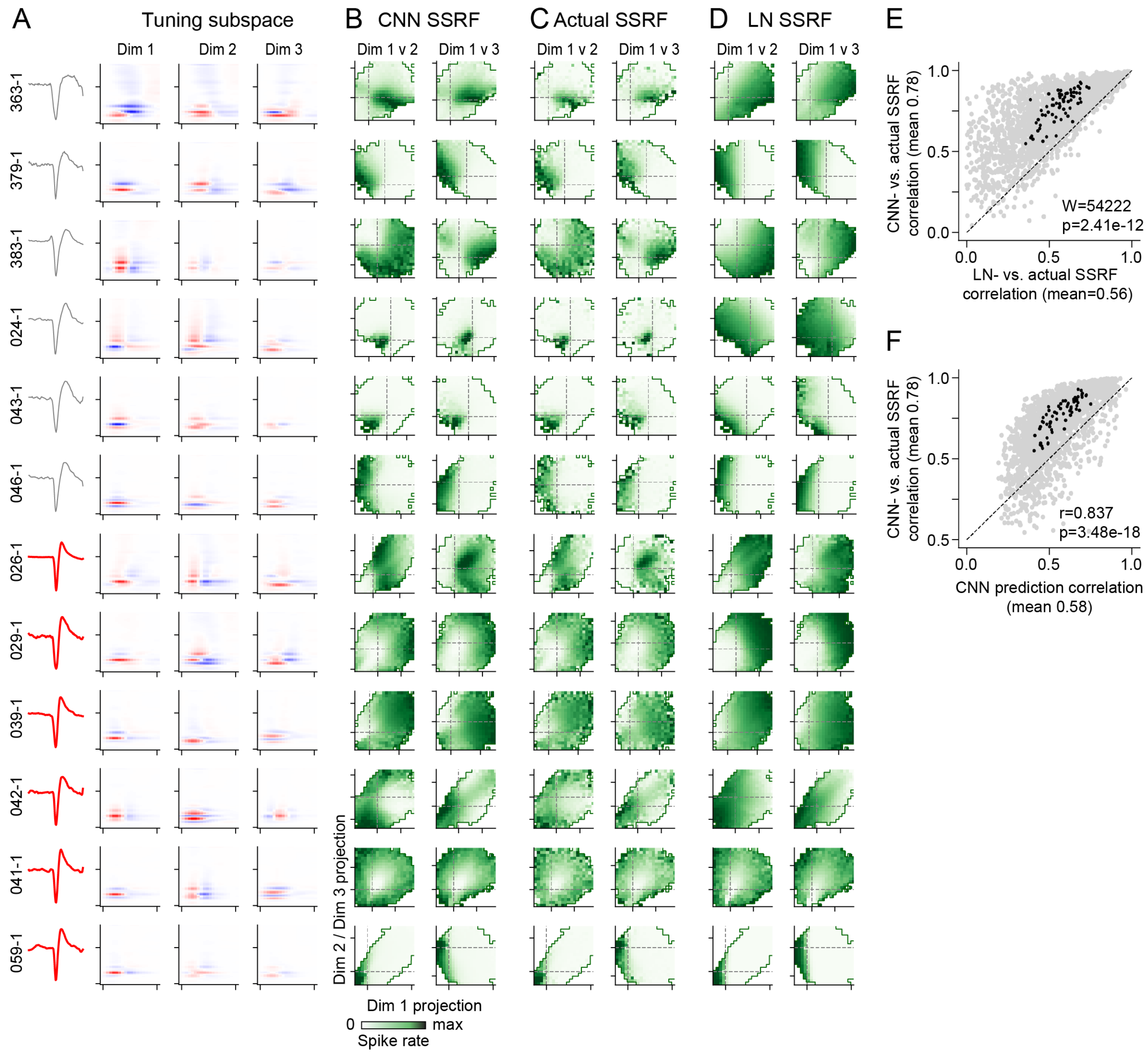


**Figure S4.** Accuracy of subspace receptive fields (SSRFs) measured from CNN encoding model. **A.** Mean spike waveforms and tuning subspace filters for 12 example units from the same recording site and plotted as in Fig. 3. **B.** Heat maps show SSRFs measured from the CNN predicted response, computed as the average spike rate for stimuli projecting to each point in the tuning subspace for dimensions 1 vs. 2 (left column) and dimensions 1 vs. 3 (right column). Darker green indicates higher spike rate. **C.** SSRFs measured from the actual PSTH response, plotted as in B. Actual SSRFs show similar features to the CNN predictions in B. **D.** SSRFs for the same neurons based on activity predicted by the LN model. These SSRFs show some correspondence to the actual SSRF but often fail to capture narrow and non-monotonic features. **E.** Scatter plot compares the correlation coefficient between the actual SSRF and LN prediction (x-axis) versus for the actual SSRF and CNN prediction (y-axis). Gray dots are individual units, and black dots show medians per recording site (n=1751 sound-responsive AC units, 67 recording sites). Correlation is consistently stronger for CNN SSRFs (p=2.4e-12, Wilcoxon signed-rank test). **F.** Scatter plot compares CNN model prediction correlation (x-axis) versus CNN SSRF correlation with actual SSRF (y-axis), for single units and site averages, plotted as in E. The accuracy with which the CNN predicts the SSRF is strongly correlated with its ability to predict the response PSTH (r=0.84, p=3.5e-18, t-test).

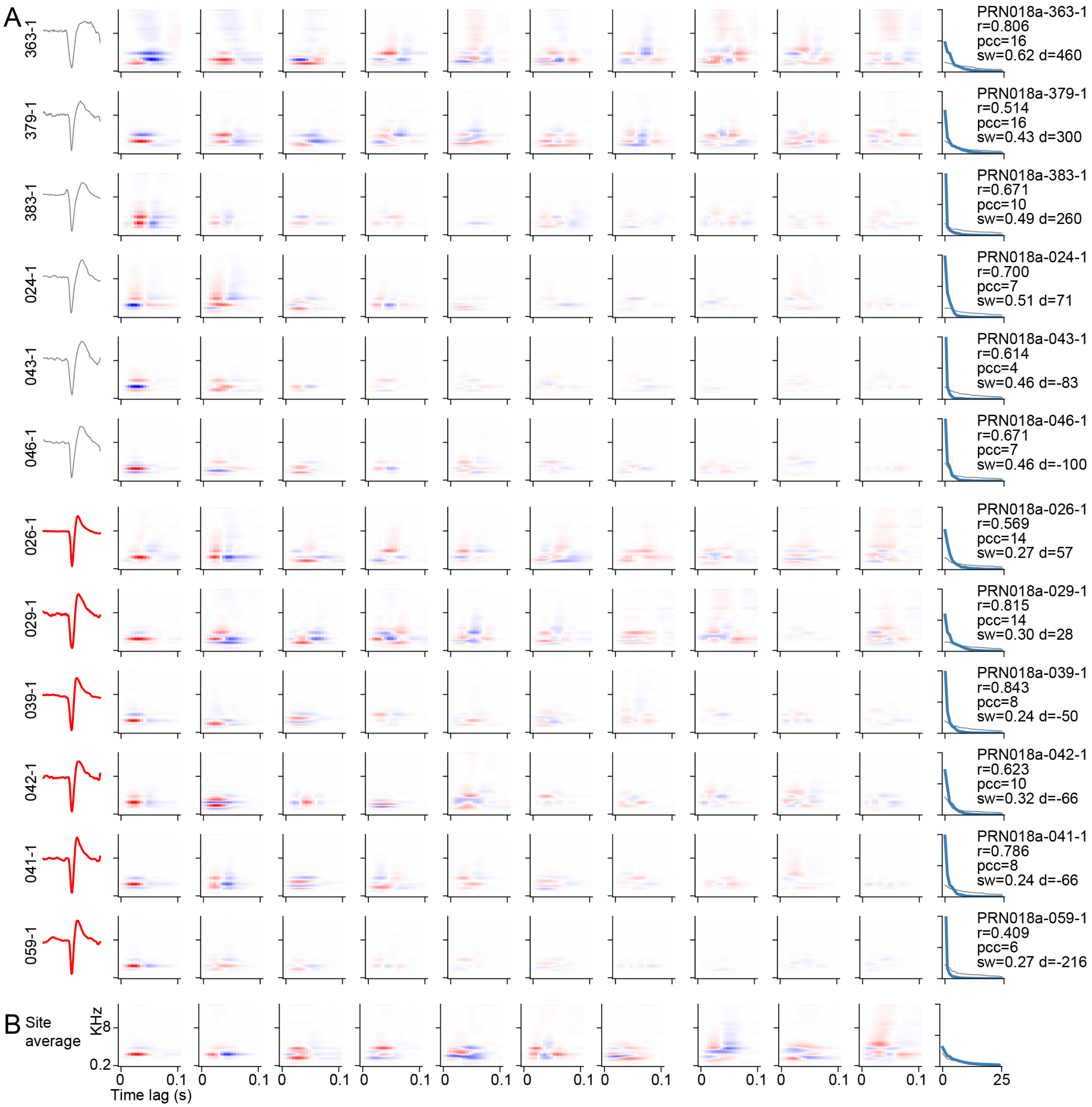


**Figure S5.** Example single-unit and recording site-wide tuning subspaces **A.** Ten highest-variance tuning subspace filters (columns) for 12 single units (rows) recorded simultaneously from the same site as in Fig. 3. The tuning subspace was computed by performing PCA across dynamic spectrotemporal receptive fields (dSTRFs) measured from the CNN encoding model fit for each unit. Left shows mean spike waveform (gray: regular-spiking; red: narrow-spiking). Right shows fraction dSTRF variance explained by each dimension. Numbers to the right indicate prediction correlation (r), number of PCs required for 95% dSTRF variance (*pcc*), spike peak-trough width in ms (sw), and depth, um below the L3/4 boundary (d). **B.** Site-wide tuning subspace filters measured for the same site by performing PCA across dSTRFs pooled across all 40 units in the site, plotted as in A.

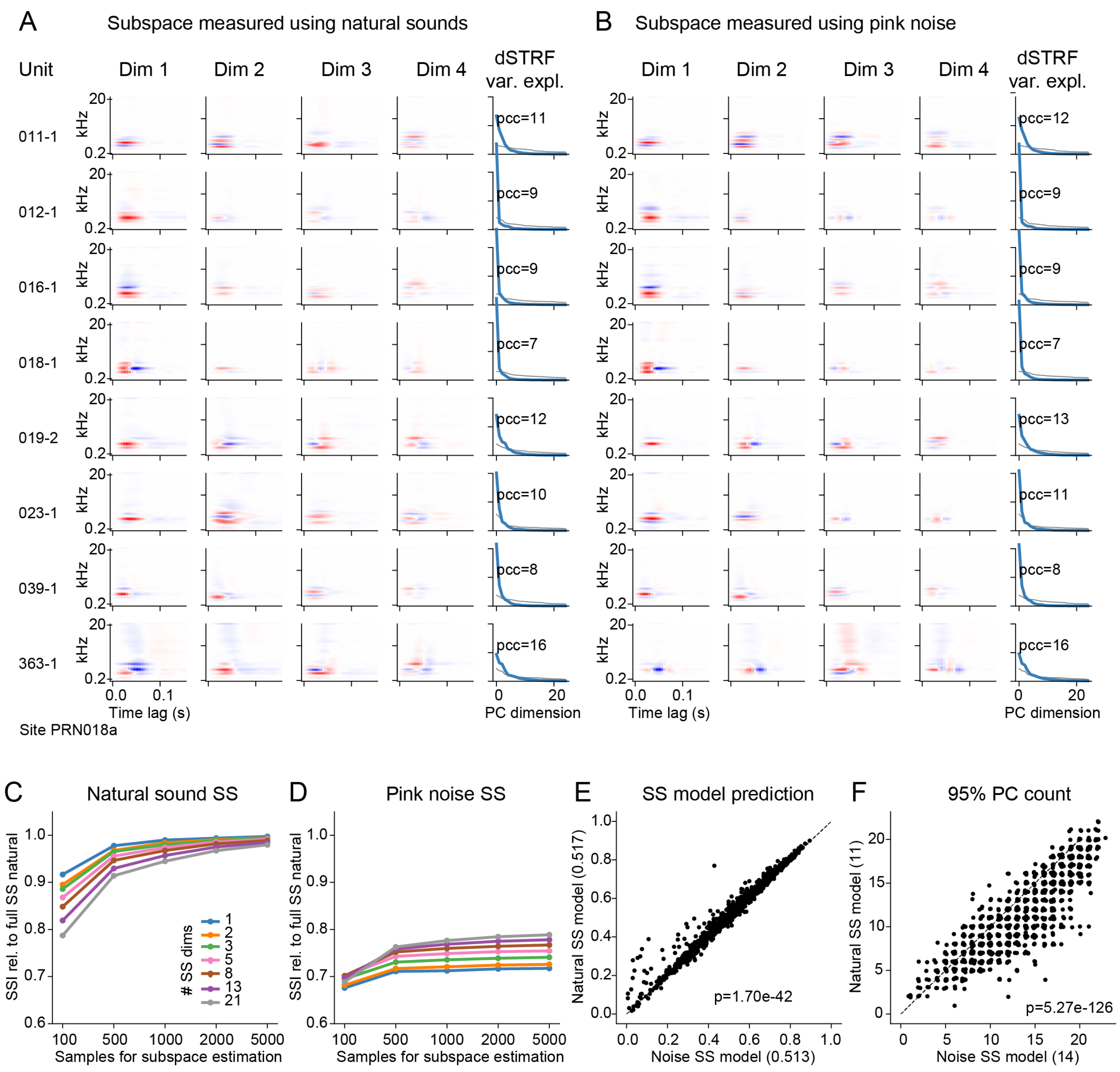


**Figure S6.** Dependence of tuning subspace on stimulus used for measurement. **A.** Examples of first four subspace dimensions measured using dSTRFs for the natural sound sequence used for training the CNN encoding model. Examples are for 8 units from the same site as in Fig. S5. **B.** Tuning subspace for the same units as in A, but measured using noise stimuli with average power spectrum matched to the original training stimuli. Subspace filters are broadly similar to the natural sound measurements in A (e.g., center frequency and modulation tuning). However, total dSTRF variance explained by the noise-based measures decreases more slowly (compare curves in right column to those in A). PC count (*pcc*) indicates the number of dimensions required to explain 95% of dSTRF variance. **C.** Dependence of subspace on the number of dSTRF samples used **for** measurement. Mean subspace similarity index (SSI) between subspace estimated with 8000 natural sound stimuli and smaller, random, non-overlapping stimulus samples, measured within-unit (n=1267 units, randomly sampling all sites in the dataset). To account for different numbers of dimensions explaining 95% of dSTRF variance, SSI was computed for varying numbers of dimensions, indicated by curves. SSI remains high, >0.95 when >2000 stimulus samples are used, indicating that subspace estimates are relatively stable. **D.** SSI between tuning subspaces measured using noise stimuli and the original natural sound subspace increase more slowly, reaching values near 0.8 for the largest set tested. **E.** Scatter plot compares model prediction accuracy for subspace encoding models fit using tuning subspaces measured using natural stimuli versus using noise stimuli. Prediction accuracy, measured using the same held-out test natural stimulus set, is similar but slightly greater for the natural stimulus subspaces (n=1267, p=1.7e-42, Wilcoxon sign test). Both models were fit using a subspace with adequate dimensions to explain 95% of dSTRF variance. **F.** Scatter plot compares number of dimensions required to explain 95% of dSTRF variance for tuning subspaces measured with natural stimuli versus noise. A small jitter is added to each point to display relative density of dots, which are constrained to integer values. Tuning subspaces measured using noise stimuli require a larger median number of dimensions (14 vs. 11, p<5.3e-126, sign test). Because uncorrelated noise stimuli span a larger space than natural sounds, they likely evoke more variable dSTRFs, thus requiring more components to explain the same amount of variance.

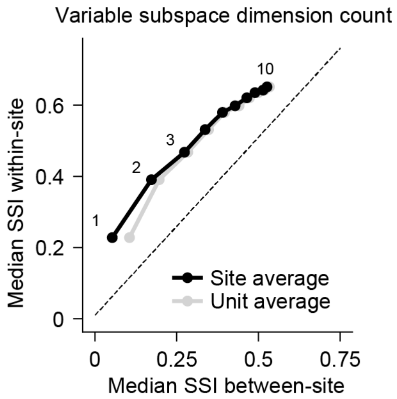


**Figure S7.** Similarity of tuning subspace within- versus between sites is independent of subspace dimensionality. Points show average subspace similarity index (SSI) between the tuning subspace for single neurons and the site-wide tuning subspace from the same recording site (within-site, computed without the current neuron) and a different recording site fit with an identical stimulus set (between-site). Numbers indicate how many subspace dimensions were included in the SSI calculation. Because subspaces were derived from population models fit using data from all the neurons, noise tended to drive higher subspace dimensions to be more similar (see Fig. S9D), both within- and between sites. Despite the trend toward increasing similarity, the within-site SSI remains consistently higher. Black dots show SSI averaged first within each site and then across all sites, and gray dots show averages across all neurons, ignoring site.

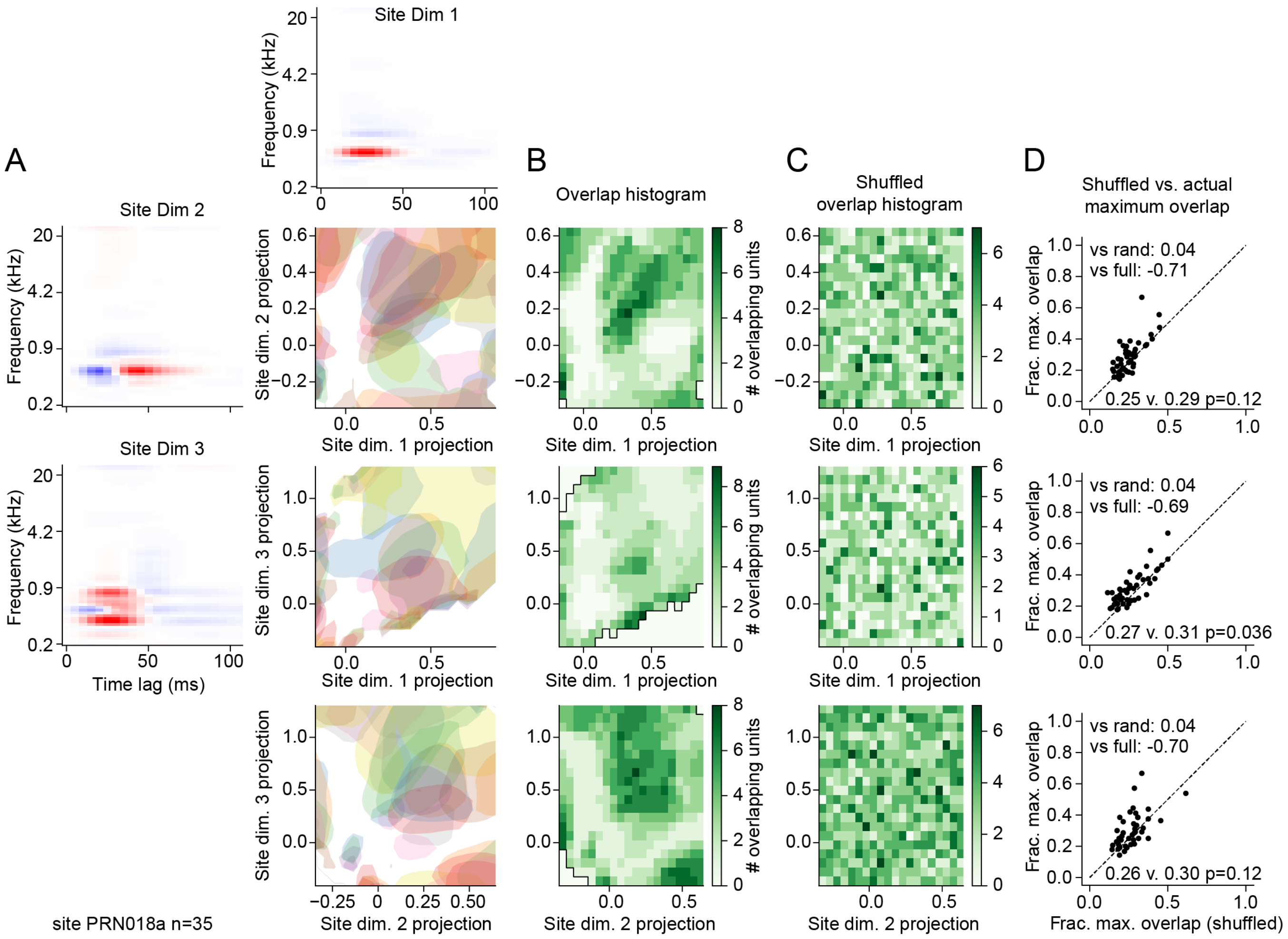


**Figure S8.** Detailed subspace tuning overlap for one recording site. **A.** Top three shared subspace filters (Site Dim 1-3) computed from aggregate dSTRF of 35 single units recorded across cortical layers in single site. Shaded plots show tuning space (contour of 80%-max response) of each unit in the shared space, projected onto pairs of subspace dimensions (top to bottom: 1 vs. 2, 1 vs. 3, 2 vs. 3). **B.** Two-dimensional histogram of overlap tuning spaces at each point in the stimulus subspace. **C.** Histogram of overlap after shuffling tuning spaces. **D.** Scatter plot shows fraction maximum overlap for actual vs. shuffled tuning spaces for each recording site (n=39 sites with >10 sound-responsive units). Numbers indicate mean difference in actual overlap from random shuffled and complete overlap (fraction=1.0), and p-values indicate significance of difference from random (Wilcoxon signed rank test).

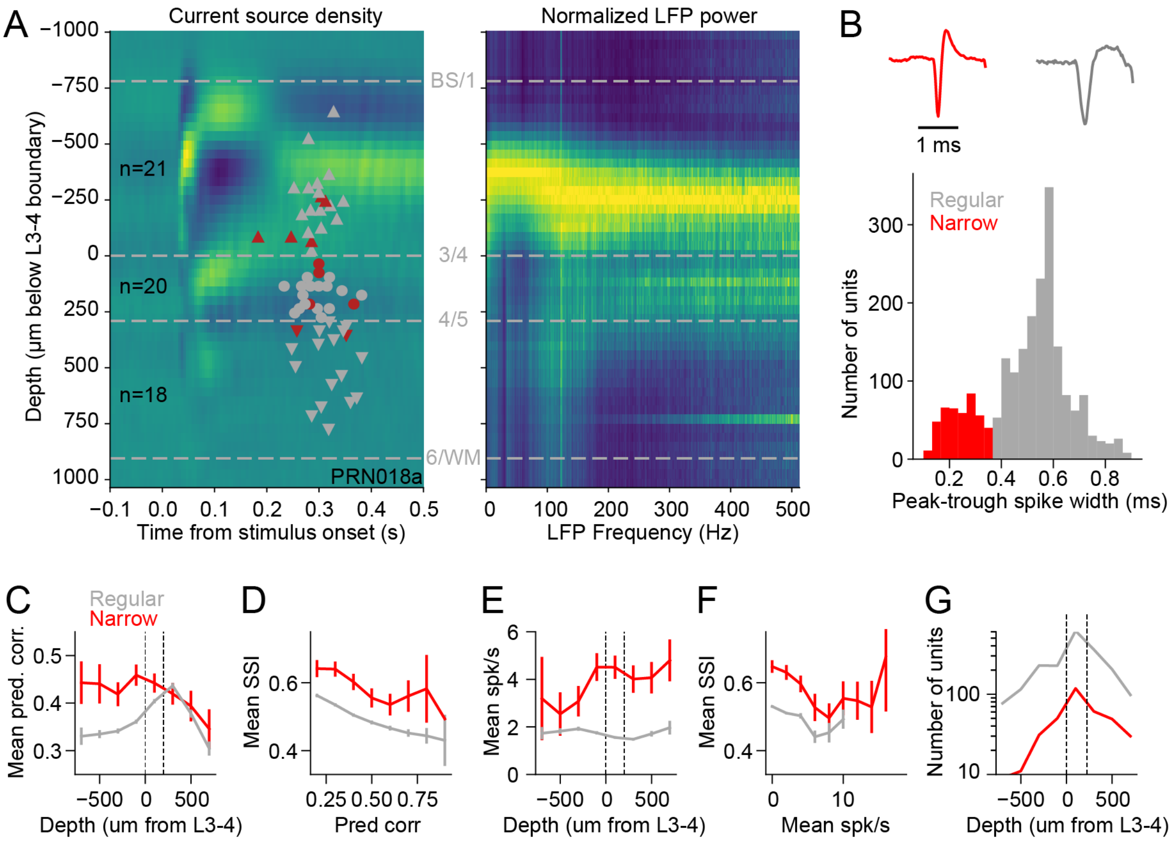


**Figure S9. A.** Current source density (CSD) evoked by narrowband noise, measured across leads of a Neuropixels probe spanning 2 mm of A1 (right). Gray/red shapes indicate depth of single regular-spiking/narrow-spiking neurons isolated during recording. Dashed lines indicate layer boundaries identified by CSD features. Local field potential (LFP) power spectrum computed across the same data, normalized to the maximum across depths. The band of high power provides a secondary indicator of L3. **B.** Histogram of spike widths, regular (R, putative excitatory)/narrow (N, putative inhibitory) classes indicated by gray/red color (n=2337 sound responsive units). **C.** Curves show mean prediction correlation by the subspace encoding model, for neurons grouped by spike width (regular or narrow, color), as a function of cortical depth. Error bars are 2 SEM. Dashed vertical lines indicate layer 3-4 and 4-5 boundaries, as measured from the LFP in panel A. **D.** Curves show mean SSI for pairs of neurons from the same recording site and with same spike width, as a function of the mean prediction correlation by the subspace encoding model. SSI tends to decrease for neurons with more accurate predictions, suggesting that low signal-to-noise biases measures of subspace tuning to be more similar. **E.** Mean overall spike rate (for each neuron over the entire natural sound dataset) as a function of spike width and cortical depth, plotted as in C. **F.** Mean SSI as a function of average spike rate for pairs of neurons, plotted as in D. **G.** Number of neurons included in SSI and TSI analyses (Figs. 5-6), as a function of spike width (color) and cortical depth (x position). Regular spiking neurons constituted approximately 80% of the sample. Larger number of neurons in middle layers (near depth 0) may reflect sampling bias, as electrode array insertions did not always span all layers of cortex.

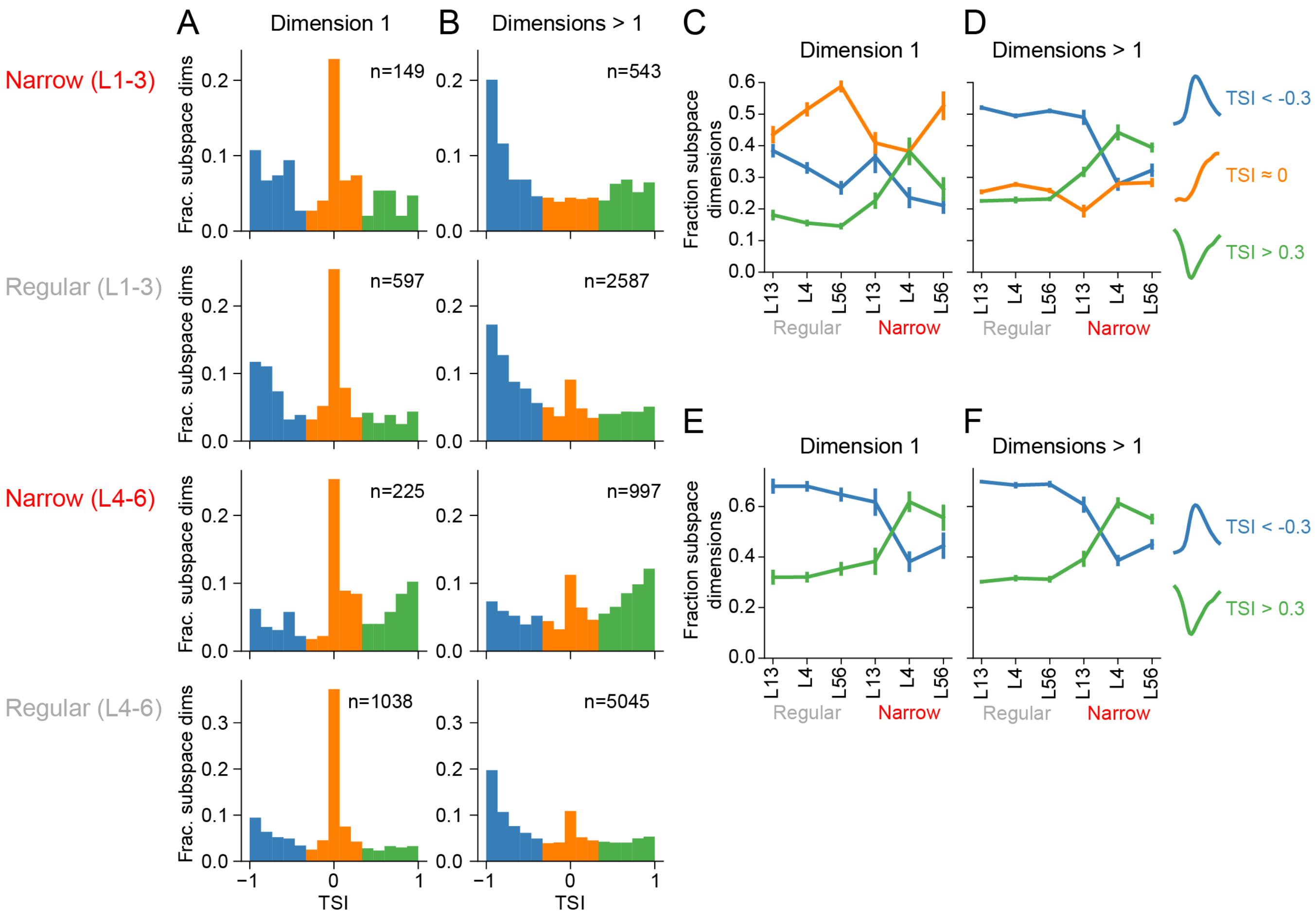


**Figure S10. A.** Histograms of TSI measured across all neurons for tuning subspace dimension 1, i.e., the dimension explaining the most variance in the dSTRF measured from the CNN-based encoding model. Data are grouped by cell type (narrow or regular spike width, deep versus superficial cortical layer). A full range of TSI values was observed, but peaks occurred at -1, 0, and 1, motivating a grouping into three sign categories, bounded at -0.3 and +0.3 (colors). For dimension 1, TSI values were commonly observed near 0, corresponding to an asymmetric nonlinearity (orange). *n* indicates the number of dimensions in each group. Data from L4 and L5-6 are pooled because they follow a similar pattern. **B.** Histogram of TSI values for all other subspace dimensions (i.e., dimensions > 1) required to explain 95% of dSTRF variance across all neurons, plotted as in A. Asymmetric nonlinearities are less common, and the frequency of negative versus positive TSI varies between cell type. Results are reported pooled across A1 and PEG (total n=1632 regular, 374 narrow, 11181 unit/subspace combinations accounting for 95% of dSTRF variance for each unit). **C.** Fraction of subspace dimension 1 in each TSI sign category (color, legend at right), grouped by spike width (regular or narrow) and cortical layer. Error bars indicate standard error on the mean, calculated by jackknifing. **D.** Fraction of subspace dimensions > 1 in each TSI sign category, plotted as in A. Across all cell types, the fraction of asymmetric TSI (-0.3 < TSI < 0.3, orange) is more common for dimension 1 than for dimensions > 1 (0.50 versus 0.26, U=1.30e7, p=1.5e-110, Mann-Whitney U-test). **E.** Fraction of subspace dimensions 1 in each TSI sign category, after excluding dimensions with asymmetric TSI, near 0 (legend at right). **F.** Fraction of smaller subspace dimensions >1 in each TSI sign category, after excluding dimensions with TSI near 0, plotted as in C.

| Parameter | Estimate | Std. Error | t value | $p(>\lvert t\rvert)$ | $\chi^2$ | $p(>\chi^2)$ |
|---|---|---|---|---|---|---|
| **A1** | | | | | | |
| (Intercept) | 0.626 | 0.0467 | 13.4 | 8.6e-4 | | |
| Spike width = narrow, regular (WNR) | -0.0474 | 0.00353 | -13.4 | < 2e-16 | 495.8 | < 2.2e-16 |
| Spike width = regular, regular (WRR) | -0.0709 | 0.00350 | -20.2 | < 2e-16 | | |
| Mean depth (D) | -0.0409 | 0.00307 | -13.3 | < 2e-16 | 566.6 | < 2.2e-16 |
| Depth difference (Δ) | -0.0230 | 0.00400 | -5.75 | 9.0e-9 | 78.32 | < 2.2e-16 |
| Mean rate (M) | -0.0138 | 0.000808 | -17.1 | < 2e-16 | 290.9 | < 2.2e-16 |
| Prediction correlation (R) | -0.0254 | 0.000783 | -32.4 | < 2e-16 | 1052.5 | < 2.2e-16 |
| WNR x D | 0.0104 | 0.00319 | 3.3 | 1.1e-3 | 52.8 | 3.5e-12 |
| WRR x D | 0.0184 | 0.00310 | 5.93 | 3.1e-9 | | |
| WNR x Δ | 0.0151 | 0.00424 | 3.55 | 3.8e-4 | 20.0 | 4.6e-5 |
| WRR x Δ | 0.0177 | 0.00409 | 4.33 | 1.5e-5 | | |
| **PEG** | | | | | | |
| (Intercept) | 0.547 | 0.0346 | 15.8 | 2.4e-4 | | |
| Spike width = narrow, regular (WNR) | -0.0395 | 0.00927 | -4.26 | 2.0e-5 | 34.1 | 4.0e-8 |
| Spike width = regular, regular (WRR) | -0.0485 | 0.00925 | -5.24 | 1.7e-7 | | |
| Mean depth (D) | -0.0107 | 0.00993 | -1.07 | 2.8e-1 | 167.4 | < 2.2e-16 |
| Depth difference (Δ) | -0.0565 | 0.00939 | -6.02 | 1.8e-9 | 475.7 | < 2.2e-16 |
| Mean rate (M) | -0.00959 | 0.00133 | -7.21 | 6.0e-13 | 51.9 | 5.8e-13 |
| Prediction correlation (R) | -0.0109 | 0.00140 | -7.81 | 6.2e-15 | 60.9 | 5.9e-15 |
| WNR x D | -0.00619 | 0.0102 | -0.61 | 5.5e-1 | 0.85 | 0.65 |
| WRR x D | -0.00396 | 0.0100 | -0.40 | 6.9e-1 | | |
| WNR x Δ | 0.0220 | 0.00969 | 2.27 | 2.3e-2 | 35.8 | 1.7e-8 |
| WRR x Δ | 0.0354 | 0.00948 | 3.73 | 1.9e-4 | | |

**Table S1.** Results of linear mixed effects regression model testing effects of neuronal cell type and depth on subspace similarity index (SSI) for pairs of neurons recorded from the same cortical site. Each pair was categorized by their constituent spike widths, either narrow-narrow (WNN)**,** narrow-regular (WNR), or regular-regular (WRR). Depth-related fixed effects for each neuron pair were average depth (D), difference in depth (Δ), and their interaction with spike width. Mean spike rate (M) and CNN prediction correlation (R) were also included as predictors, and animal was included as a random effect. Results are reported separately for A1 (n=1459 regular, 205 narrow, 42140 pairs) and PEG (n=532 regular, 64 narrow, 18833 pairs). T values indicate effect size for individual main effects and interactions. Type II Wald $\chi^2$ tests assess the overall significance of each predictor variable. Model fitting and statistical analysis performed in R.

| Parameter | Estimate | Std. Error | z value | p(>\|z\|) | $\chi^2$ | p(>$\chi^2$) |
|---|---|---|---|---|---|---|
| **A1** | | | | | | |
| (Intercept) | -0.39 | 0.17 | -2.31 | 0.0209 | | |
| Spike width = regular (WR) | -0.34 | 0.13 | -2.69 | 0.0073 | 62.00 | 3.4e-15 |
| Depth = L4 (DL4) | 0.64 | 0.16 | 4.10 | 4.1e-05 | 3.89 | 0.143 |
| Depth = L45 (DL56) | 0.36 | 0.16 | 2.28 | 0.0224 | | |
| Mean rate (M) | 0.22 | 0.03 | 6.34 | 2.2e-10 | 40.25 | 2.2e-10 |
| Prediction correlation (R) | -0.08 | 0.03 | -2.43 | 0.01496 | 5.92 | 0.0150 |
| WR x DL4 | -0.66 | 0.18 | -3.68 | 2.3e-4 | 13.57 | 0.00113 |
| WR x DL56 | -0.33 | 0.18 | -1.81 | 0.0696 | | |
| **PEG** | | | | | | |
| (Intercept) | -1.03 | 0.30 | -3.49 | 4.8e-4 | | |
| Spike width = regular (WR) | 0.34 | 0.24 | 1.43 | 0.1519 | 5.29 | 0.0214 |
| Depth = L4 (DL4) | 1.23 | 0.29 | 4.24 | 2.2e-5 | 11.95 | 0.00254 |
| Depth = L45 (DL56) | 1.35 | 0.32 | 4.26 | 2.1e-5 | | |
| Mean rate (M) | 0.26 | 0.06 | 4.28 | 1.9e-5 | 18.33 | 1.8e-05 |
| Prediction correlation (R) | -0.18 | 0.05 | -3.28 | 0.00102 | 10.78 | 0.00102 |
| WR x DL4 | -1.04 | 0.31 | -3.32 | 9.0e-4 | 14.75 | 6.3e-4 |
| WR x DL56 | -1.15 | 0.34 | -3.38 | 7.2e-4 | | |

**Table S2.** Results of generalized linear mixed effects regression model testing effects of neuronal cell type and depth on tuning symmetry index (TSI: whether positive or negative) for each neuron. Mean spike rate (M) and CNN prediction correlation (R) were also included as predictors, and animal was included as a random effect. Results are reported separately for A1 (n=1246 regular, 310 narrow, 5644 unit/subspace combinations) and PEG (n=386 regular, 64 narrow, 2129 unit/subspace combinations). z values indicate effect size for individual main effects and interactions. Type II Wald $\chi^2$ tests assess the overall significance of each predictor variable. Model fitting and statistical analysis performed in R.
